# BaseQTL: a Bayesian method to detect eQTLs from RNA-seq data with or without genotypes

**DOI:** 10.1101/2020.07.16.203851

**Authors:** Elena Vigorito, Wei-Yu Lin, Colin Starr, Paul DW Kirk, Simon R White, Chris Wallace

## Abstract

Available methods to detect molecular quantitative trait loci (QTL) require study individuals to be genotyped. Here, we describe BaseQTL, a Bayesian method that exploits allele-specific expression to map molecular QTL from sequencing reads even when no genotypes are available. When used with genotypes, BaseQTL has lower error rates and increased power compared with existing QTL mapping methods. Running without genotypes limits how many tests can be performed, but due to the proximity of QTL variants to gene bodies, the 2.8% of variants within a 100kB-window that could be tested, contained 26% of QTL variants detectable with genotypes. eQTL effect estimates were invariably consistent between analyses performed with and without genotypes. Often, sequencing data may be generated in absence of genotypes on patients and controls in differential expression studies, and we identified an apparent psoriasis-specific effect for *GSTP1* in one such dataset, providing new insights into disease-dependent gene regulation.

## Introduction

Genome-wide association studies (GWAS) have identified thousands of genetic variants associated with human disease, but their individual mechanisms remain largely unknown. The majority of variants are located outside coding regions, and are presumed to have regulatory function^1^. Thus mapping these variants to their direct target gene(s) is instrumental to understand the molecular mechanisms that predispose to disease. Direct regulatory effects result when the genetic variant and the target gene are located on the same chromosome, typically less than 1 MB apart, and the genetic variant only affects the expression of the gene copy on its same chromosome. Common biological processes regulated by such variants include transcription, RNA stability or splicing. These variants are commonly referred as cis-QTLs, for example cis-eQTL for those which regulate expression.

Mapping the causal genes should be possible by testing whether these variants have an effect on gene expression and large studies have been established to map regulatory variants across a diverse range of tissues. Nonetheless, multiple studies have failed to enumerate more than a minority of genes regulated by disease-associated variants^2–4^. This is likely due to the highly context-specific effects of genetic variants on gene expression^5,6^ and that most eQTL studies to date have focused on healthy individuals and bulk tissues ^7^. Gene expression data from specific cell types in disease contexts appear to be more informative for interpreting disease-associated variants,^8,9^ but such datasets have often been generated in the context of biomarker studies or in efforts to understand the disease process rather than the genetics of gene expression. Therefore such datasets are commonly small (<100 samples), may be designed to compare the same tissue in different contexts (e.g. disease activity) and/or may have no genotype data available.

Standard eQTL studies estimate average fold change in expression according to allelic dose by comparing expression between genotyped individuals. eQTL methods are typically embedded within broader statistical analysis environments, so that custom analyses comparing fold change estimates between sample contexts can be explored. Power can be improved by approaches which additionally exploit imbalance in gene expression between chromosomes within heterozygous individuals, so-called allele specific expression (ASE). However, ASE software is generally limited to detecting eQTLs so that it is difficult to extend to related questions such as comparing effects between conditions. To the best of our knowledge, all ASE methods to date require study subjects to be genotyped.

Here, we propose a new method, BaseQTL, for ASE analysis. By adopting a Bayesian approach, we can incorporate information from existing large eQTL studies which allow us to shrink extreme fold change estimates and improve accuracy in a way analogous to moderation of variance estimates in differential expression analyses ^10^. We embed this within a standard Hamiltonian Monte Carlo (HMC) environment ^11^, allowing researchers to develop flexible analytic approaches appropriate to their data. The phase of regulatory SNPs and the genic SNPs at which allelic imbalance is measured is unknown, and standard ASE methods infer phase from the individuals within each study. Using a Bayesian approach, we also exploit external reference genotype panel data to improve phasing accuracy. Our model treats phase as latent (unknown), and we extend this latent structure to also treat candidate regulatory SNP (cis-SNP) genotype as unknown, allowing us to analyse studies without separate genotype data.

We used 86 LCL samples from GEUVADIS^12^ for whom genotypes and RNA-seq data are publicly available, to benchmark BaseQTL against standard eQTL and ASE methods and to compare the results of BaseQTL run with genotypes either available or masked.

We then used our method to call eQTLs in a publicly available RNA-seq data from 94 psoriasis and 82 normal skin samples^13^. The ability to call eQTLs in existing patient RNA-seq datasets, even when no genotypes are available, may help us better understand the mechanism underlying established GWAS signals for complex diseases.

## Results

### Basic model to detect cis-eQTL

To detect cis-QTL using RNA-seq data, standard methods test the association between a genotyped variant within a specific distance of a genome feature (gene, ChIP-peak, etc) and the total count of short reads mapped to the feature. ASE models additionally exploit the knowledge that, if the cis-SNP was associated with expression, we would expect this to result in imbalanced expression between the two chromosomes in individuals heterozygous at the cis-SNP. Phase-aware cis-QTL methods such as RASQUAL^14^, WASP^15^ or TReCASE^16,17^ substantially improve the power of sequencing-based QTL mapping by jointly modelling the differences in total read counts mapping to the feature *between* individuals, and the allelic imbalance at phased heterozygous feature SNPs (fSNPs) *within* individuals, as functions of the genotype at the candidate cis-SNP (Fig. 1a). RASQUAL models total gene counts with a negative binomial (NB) distribution, WASP with a beta negative binomial and TReCASE with a Poisson-NB mixture. For allele-specific signals, RASQUAL, WASP and TReCASE all use a beta-binomial model, with some differences, compared in ^14^. We begin by describing the TReCASE model^16^; ^17^ which expresses the likelihood as a product of between- and within-individual components.

**Figure 1.**
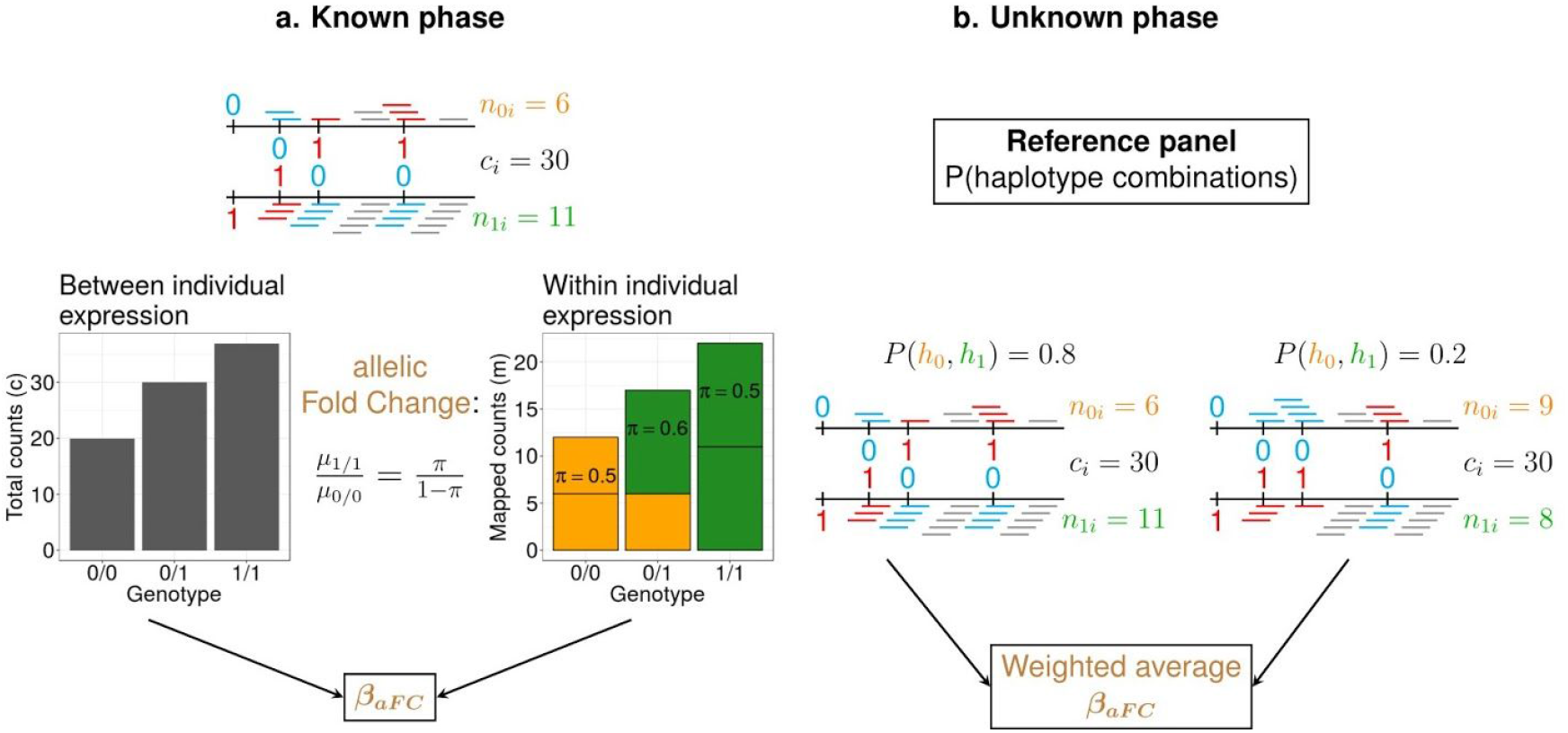
Schematic representation of BaseQTL with observed genotypes. RNA-seq reads overlapping the reference or the alternative allele for a SNP are depicted in blue or red, respectively; gray reads do not overlap SNPs. (a) Joint model combining between and within individual variation when genotypes are observed and phase known. The top panel illustrates the “true” haplotype pair formed by a cis-SNP and 3 fSNPs within a gene in a heterozygous individual for the cis-SNP. Allele specific expression (ASE) is measured as the proportion of reads mapping fSNPs within the haplotype carrying the cis-SNP alternative allele (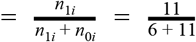). The total counts mapped to the gene are indicated by c_i._. The lower panel shows how between and within individual expression are connected for cis-eQTLs. μ_0/0_ and μ_1/1_ correspond to the expected total reads (*c*_*i*_) in homozygous individuals for the reference or alternative allele respectively, whereas π corresponds to the expected mean for the ASE proportion. The model estimates β_aFC_=μ_1/1_ / μ_0/0_. (b) We extend this model to account for unknown phase by assuming that β_aFC_ follows a mixture distribution conditional on phase, which is treated as a latent variable. Phase probabilities are estimated from a large external reference panel conditional on the observed genotypes.

Between-individual signals were modelled by negative binomial regression of the total read counts on the genotype of the cis-SNP (*G*_*i*_ = 0,1,2) allowing for additional covariates (library size, principal component loadings, etc.). This is a standard eQTL model. TReCASE assumes that the phase at the cis-SNP and any heterozygous fSNPs is known, so that the number of reads mapping to the haplotype carrying the alternative allele for the cis-SNP, out of the total number of ASE mapped reads can be modelled using a beta binomial distribution. Between and within individual components are connected by a single parameter β_*aFC*_, which corresponds to the expected log-allelic fold change (aFC) between individuals homozygous for the alternative and reference alleles at the tested cis-SNP (**Fig. 1a** and Online methods). We use the same model and introduce a series of novel extensions which are described in the coming sections. We assessed each extension using a modest sample size of 86 RNA-seq lymphoblastoid cell lines (LCLs) from European individuals (GBR) generated by the GEUVADIS project, for which genotypes are available.^12^ This sample size was chosen to match the anticipated scale of real-world datasets where samples < 100 are common. To limit computational load, we restricted our analysis to all 264 expressed genes on chromosome 22 and cis-SNPs within 1 MB of each gene, thinned to r^2^<0.9.

### Accommodating unknown phase

With short read data, phase cannot be known with certainty and our first extension was to treat phase as latent. We replaced the within-individual component of the TReCASE likelihood by a sum of beta binomial contributions conditional on haplotype phase, weighted by their respective probabilities conditional on the unphased genotypes at the fSNPs and cis-SNPs (**Fig. 1b** and Online methods). We calculate these probabilities from 5008 phased haplotypes from the cosmopolitan 1000 Genomes phase-3 reference panel to calculate possible phased haplotype pairs and their relative probabilities. Consistent with previous reports that cosmopolitan reference haplotype panels are preferred,^18^ we found that our method is generally robust to perturbations in the reference panel but, when mismatches between samples and reference panel are extreme, there is loss of power rather than any increase in false positives (**Supplementary Table 1**).

### Modelling reference sequence mapping bias

Reference sequence mapping bias - the tendency of reads to map more easily to the reference sequence allele - can cause allelic imbalance to be overestimated in favour of the reference allele, and hence false positive ASE results ^19;20; 21; 15^. As expected, raw estimates of allelic imbalance in our GEUVADIS data subset were indeed skewed towards over-representation of the reference allele, consistent with reference mapping bias (**Fig. 2b and Supplementary Fig. 4c**).

**Figure 2.**
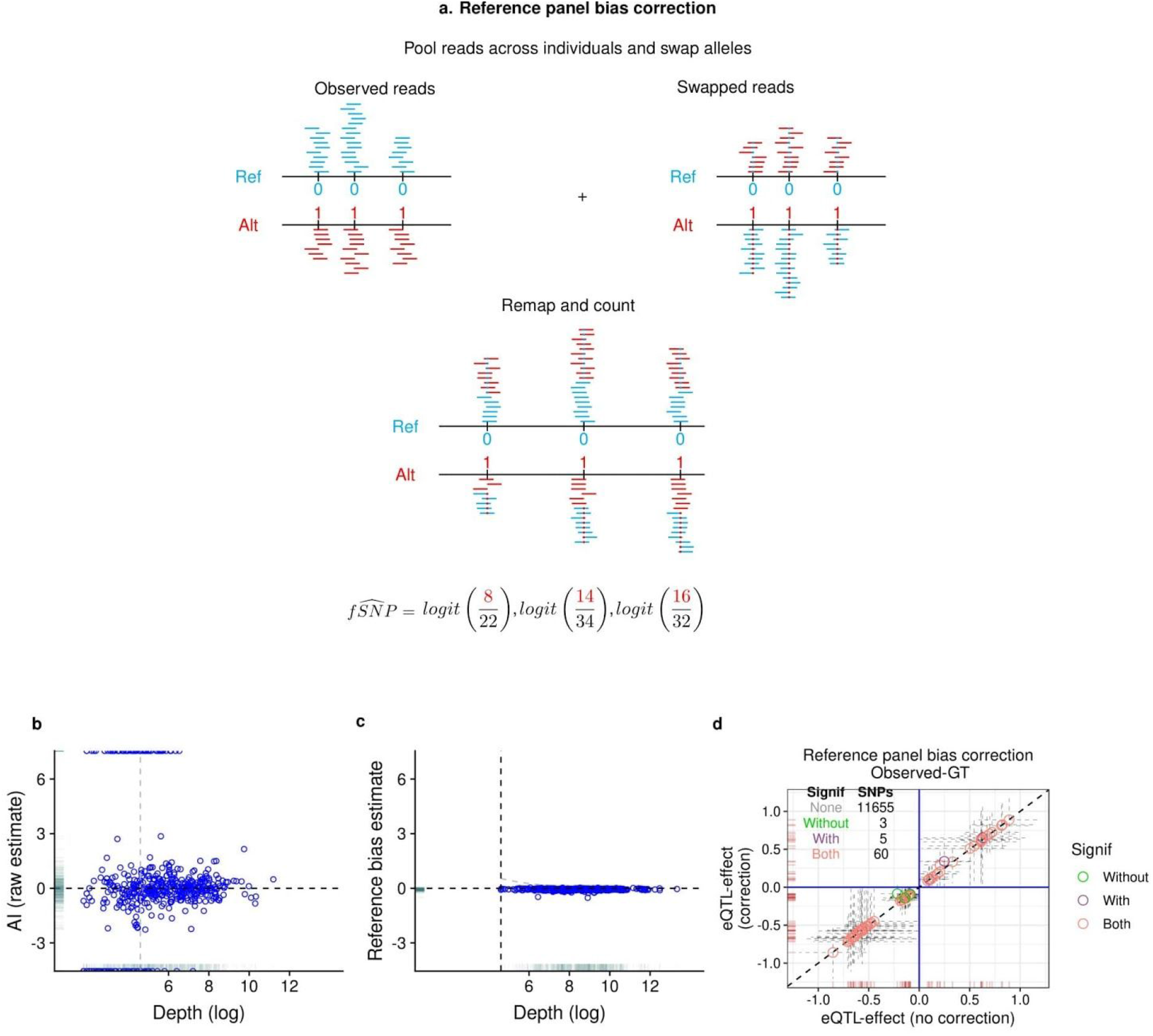
Reference mapping bias correction (a) Schematic representation of our method to correct for reference panel bias. For each read that maps to an fSNP we create a new read in which the allele of the fSNP is swapped (represented as a blue dot in a red read (alt -> ref) or a red dot in a blue read (ref -> alt). The pooled reads, which have a 50:50 ratio of reads carrying the reference or alternative alleles at each fSNP, are remapped, and the number of reads mapping to each allele stored. (b). For each fSNP we calculated the proportion of reads overlapping the alternative allele across all heterozygous individuals, which we refer to as the raw estimate of allelic imbalance (AI). The plot shows logit transformed raw estimates for AI against depth for each fSNP; mean AI is −0.05 and appears unrelated to depth. The horizontal line indicates no allelic imbalance, the gray vertical line is displayed to ease comparison with c and d. (c) Logit AI estimates obtained as described in a, have a similar mean to raw estimates (−0.06), but much smaller variance. The vertical line indicates the read threshold selected for including estimates for inference (minimum 100 reads across all samples, Online Methods). (d) The effect of reference panel bias correction is a small attenuation in effect estimates. Each symbol corresponds to a gene-SNP association comparing the eQTL estimates (log_2_ allelic fold-change) obtained with or without applying our reference bias panel correction.

Previous approaches to mitigate this phenomenon remove reads with evidence of mapping bias, while recognising that discarding data is expected to reduce power^15^. Instead of discarding reads, we model bias explicitly using a random intercept per fSNP. We modified the procedure used by WASP^15^ to identify reads susceptible to mapping bias. Reads are mapped to the genome to identify reads overlapping known SNPs. For each such read, we generate a new pseudo read in which the observed alleles in the read are swapped to the unobserved allele of the SNPs (**Fig. 2a**). The union of these observed and pseudo reads have exactly equal representation of reference and alternative alleles for each fSNP. We favoured this approach over simulating reads as we expect to have a more accurate representation of the variables that affect mappability such as genotype errors, base quality or number of polymorphisms per read.

We realigned this union of reads, and found estimated allelic imbalances showed similar skew towards the reference allele, but with much greater consistency between SNPs (**Fig. 2c**). This is due to leveraging many more reads compared to raw estimates (twice the total mapped reads compared to the subset of reads mapped to heterozygous individuals) as well as removing random noise due to true ASE (**Fig. 1a, 2b and c**). We used these estimates of allelic imbalance in these union of reads to define the parameters of a prior distribution for the random intercept, and found that adjusting for estimated reference mapping bias this way resulted in a small attenuation of eQTL effect estimates, with log fold changes on average 0.3% smaller (**Fig. 2d**).

### Levering information on eQTL effect sizes in external datasets

We leveraged fold change estimates from a study of Epstain-Barr virus transformed lymphocytes with 147 individuals from GTEx^22^ to train our prior on β_*aFC*_ by fitting a Gaussian mixture model (see Online Methods). This identified a mixture of a narrow distribution (97%, sd=0.03) and a broader distribution (3%, sd=0.35), both centered on 0 (**Supplementary Fig.1**). Similar results were obtained when we used GTEx blood or skin samples (**Supplementary Table 2)**. This informed prior shrunk 99.7% of β_*aFC*_ estimates under an uninformed prior towards 0 whilst preserving a strong correlation (rho=0.98, p<10^−16^) between eQTL effects at signals that were significant under the informed prior (**Supplementary Fig.1**). Moreover, the positive predictive value (PPV, proportion of significant hits detected by each method also detected in the gold standard), measured against a “gold standard” of a published list generated by conventional eQTL analysis called at 1% FDR from 462 GEUVADIS individuals,^23^ increased from 0.25 to 0.9.

### Benchmarking of BaseQTL against standard methods

We benchmarked BaseQTL against two other methods: standard linear regression, widely used in eQTL analysis, and RASQUAL^14^ representative of methods that jointly model between and within individual variation in a frequentist framework. As BaseQTL shrinks eQTL effects via a prior distribution, we also run BaseQTL modelling between individual variation only (negative binomial distribution) to disentangle the effect of the prior from the ASE modelling. For the same gene-SNP associations we compared eQTL calls by the four methods against the “gold standard” analysis of 462 GEUVADIS individuals.^23^ For each method we used a range of significance thresholds (Online Methods) to calculate the “sensitivity” (proportion of gold standard hits detected by each method) and the PPV for eQTLs or eGenes. We selected the same samples and genes as in the previous section but decreased the cis-window to 100kB as 54/264 genes could not be run by RASQUAL within a 1MB window. Using the smaller cis-window only 5 genes failed with RASQUAL. For the same gene-SNP associations (35,083 over 259 genes), BaseQTL outperformed both other methods (**Fig. 3a,c**) achieving the highest PPV and sensitivity trade-off. For example, for a 0.1% FDR, BaseQTL called 23 eGenes of which 22 were also called in the gold standard, linear model 14 out of 14 and RASQUAL 36 of which 22 were called by the gold standard. Expanding the cis-window to 1 MB allowed us to test 199,563 gene-SNP associations over 264 genes, showing a similar trend with BaseQTL outperforming the linear model (**Fig. 3b,d**). Interestingly, the improvement of BaseQTL over other methods appears to be due to ASE modelling as running BaseQTL only modelling between individual variation had similar performance to the linear model (Fig. 3).

From now on we refer to significant associations to those in which 0 was excluded from the posterior 99% credible interval, which corresponded to the highest PPV for BaseQTL in **Fig 3**. Overall, using BaseQTL on chromosome 22 (264 genes) we detected 192 eQTLs associated with 30 genes. Of those, 172 (90%) were replicated in the analysis of our gold standard GEUVADIS dataset corresponding to 24 eGenes (80%).

**Figure 3.**
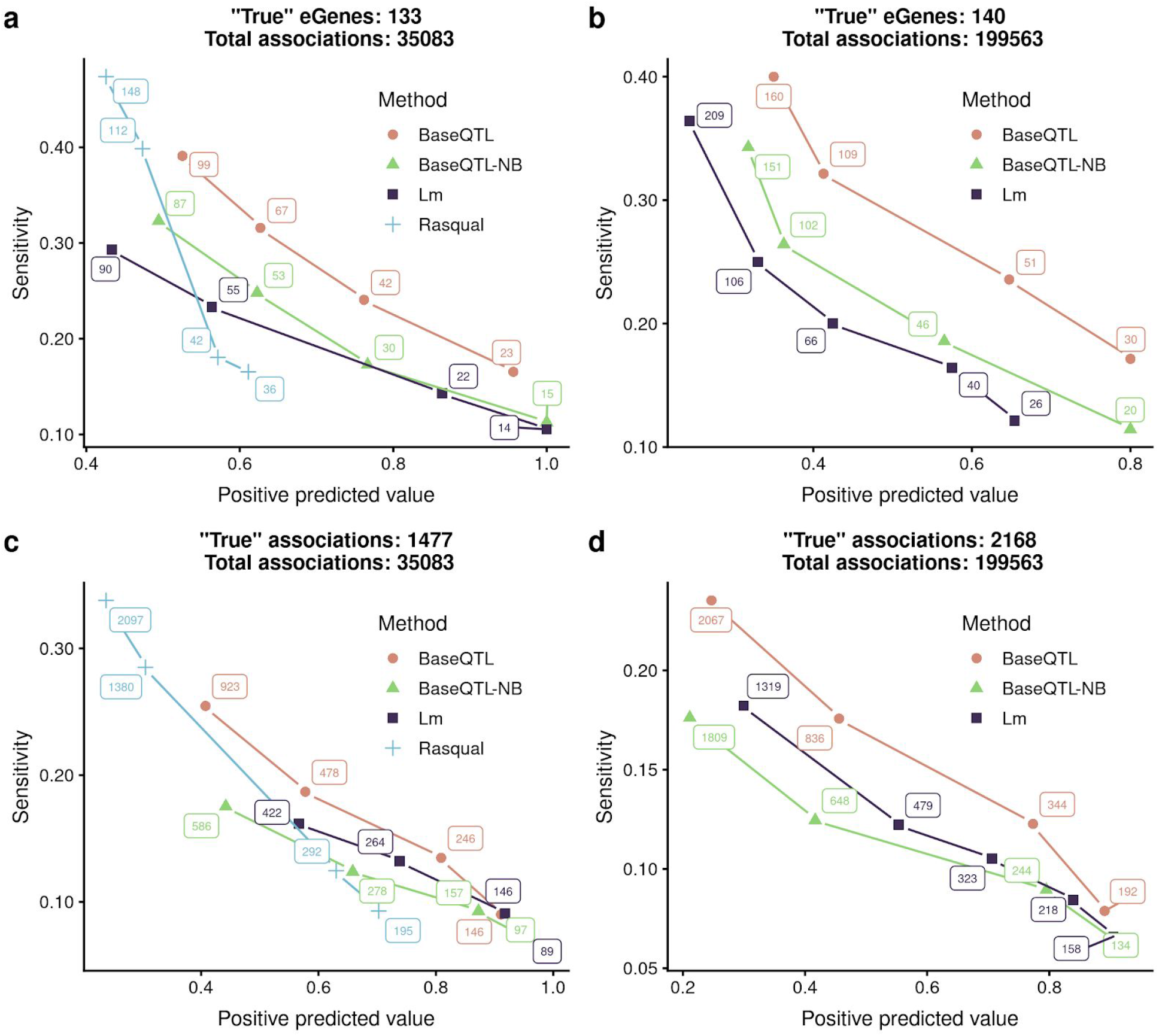
Benchmarking BaseQTL with observed genotypes using published analysis of all 462 individuals from GEUVADIS dataset as a gold standard. Analysis was performed with BaseQTL, BaseQTL modelling between individual signals only (BaseQTL-NB), linear model and when possible with RASQUAL using a sub-samples of 86 individuals from the GEUVADIS project. For each method, significant eQTLs were called for a range of FDR and positive predictive value and sensitivity were calculated relative to the gold standard. In addition, we calculated the number of eGenes called by each method by counting the number of genes with at least one significant association. The total number of significant associations or eGenes are shown at each point. **a,c** A total of 35,083 cis-SNPs within 100kB of 259 genes on chromosome 22 were analysed with the indicated method, of which 1477 (133 eGenes) were significant in the gold standard. **b,d** As in (a,c) except that a wider cis-window of 1MB within 264 genes in chromosome 22 was used. RASQUAL was excluded from the analysis as 54 genes failed to run due to the larger numbers of regulatory SNPs. Under these conditions we analysed 199,563 gene-SNP associations of whom 2168 were significant in the gold standard corresponding to 140 eGenes.

### Detecting cis-eQTLs in datasets with no genotypes

To the best of our knowledge, all ASE models to date require genotypes for the candidate cis-SNP and the fSNPs to be known, but this prevents their use in many patient datasets originally collected for other purposes. BaseQTL can be used for RNA-seq only datasets. We called fSNP genotypes by mapping feature reads to the reference genome and extended our latent model structure to treat the genotype at the candidate cis-SNP as latent, inferred probabilistically from the fSNP genotypes at the same time as haplotypes (**Fig. 1b** and Online Methods). For simplicity, we shall refer to “observed genotypes” when genotypes are measured by DNA-sequencing, and “hidden genotypes” when we only use RNA-seq for mapping eQTLs.

Genotyping error could have a major impact if not adequately controlled. In particular, homozygous fSNPs miscalled heterozygous may lead to false positives because those mis-typed fSNPs will show strong allelic imbalance. We took multiple approaches to controlling genotyping errors (**Supplementary Note**). Of the 498 fSNPs called in the 86 samples by DNA-seq (42,828 calls), 219 were excluded, and we were able to call 17,268 genotypes over 279 fSNPs with a 0.7% mismatch error (**Supplementary Fig. 2–3, Supplementary Table 3 and Supplementary Note).** While small, the mismatch rate is slightly higher than the error rate reported for short-read DNA sequencing, 0.1% to 0.6%, depending on the platform and the depth of coverage ^24^.

We assessed the impact of using RNA-seq to call fSNPs by running BaseQTL with hidden genotypes for the cis-SNPs and the same fSNPs either genotyped by RNA-seq (RNA-fSNP) or by DNA sequencing (DNA-fSNP). Estimated effects were strongly correlated (rho=0.89, **Supplementary Fig. 4a**), and all the eGenes detected with RNA-fSNPs were also called using DNA-fSNPs. We expect loss of power due to missing calls but overall these results indicate that our method is robust to genotyping errors from RNA-seq.

Next, we examined the effect of cis-SNP imputation by running BaseQTL with observed genotypes for fSNPs and with observed or hidden genotypes for the cis-SNP. We use a standard measure to assess imputation quality (Online Methods) and limited the analysis to cis-SNPs with imputation r^2^≥0.5. The imputation of the cis-SNP produces more variability on the eQTL effects than genotype errors on the fSNPs (**Supplementary Fig. 4b**).

Finally, we conducted parallel analyses by BaseQTL either with DNA-seq genotypes or RNA-seq only. Of the 264 genes on chromosome 22 tested with observed genotypes, we were able to impute genotypes for 75 (28%). We selected a smaller cis-window than before (100 kb) becuase our method for hidden genotypes strongly relies on accurate haplotype phasing, which decreases with distance. Further quality control aiming to reduce false positive rate by examining the consistency of the ASE signal across fSNPs for those genes with significant associations (Online Methods) excluded 2 genes as possible false positives. Thus, within 100 kb, we were able to assess only 1,257 gene-SNP associations with hidden genotypes compared to ~45,000 with observed genotypes (2.8%). However, both testable gene-SNP pairs with hidden genotypes and significant associations seen with observed genotypes tended to be closer to genes (**Fig. 4a**), in line with previous reports ^25^, so that the proportion of significant associations discovered with hidden genotypes (40 significant out of 1,257 [3%]) was ten-fold that with observed genotypes (153 significant out of 45,000 [0.3%]).

**Figure 4.**
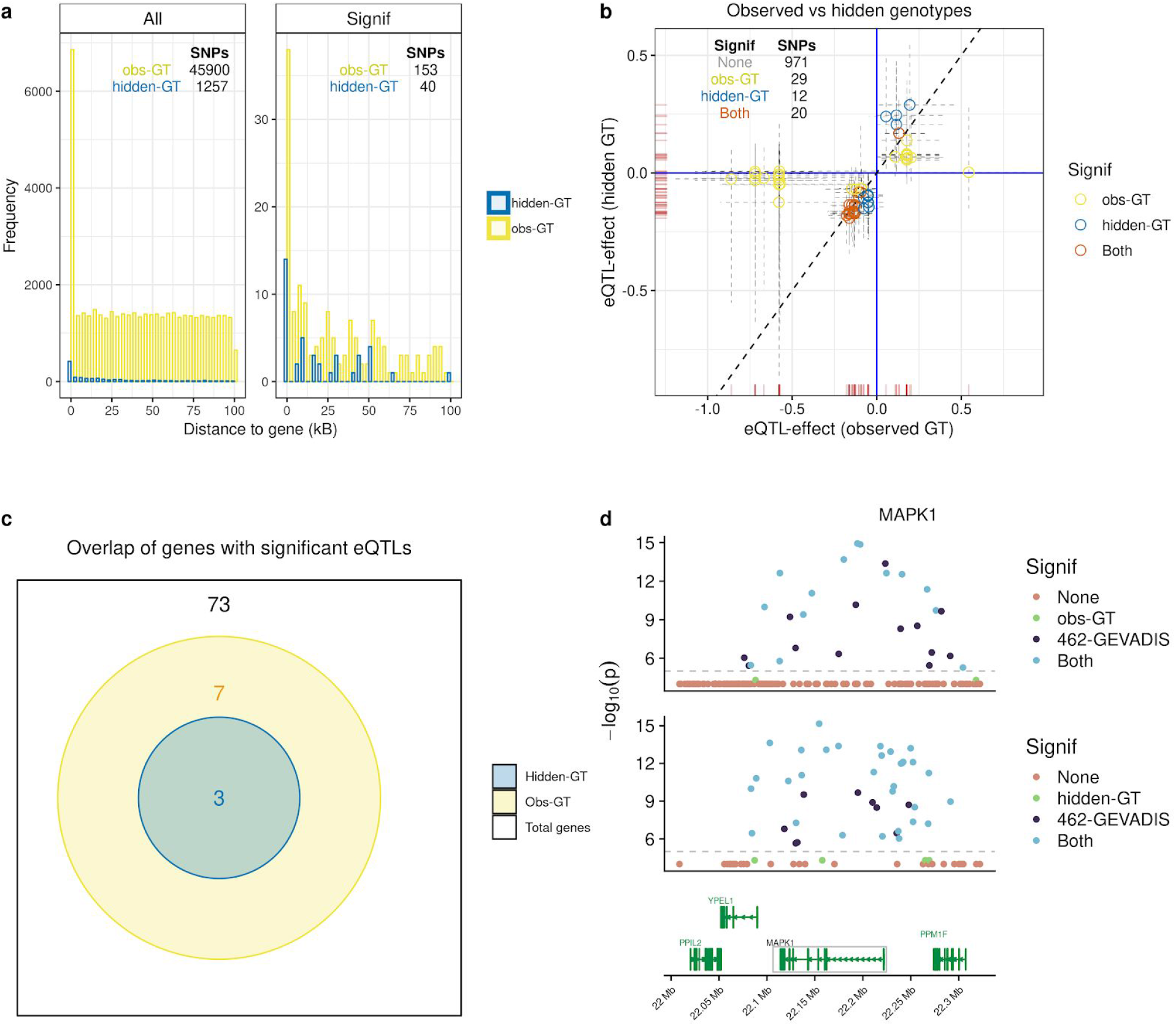
eQTL effects estimated with BaseQTL with observed or hidden genotypes. With hidden genotypes only cis-SNPs with imputation score ≥0.5 were tested. (a) For each cis-SNP the distance to the gene was calculated: for those upstream of the gene, as distance to the start of the gene; for those dowsntream of the gene, as distance to the end of the gene and for those within the gene, as 0. cis-SNPs tested with hidden hidden genotypes (blue) were concentrated closer to genes than observed genotypes (yellow), but the loss of information is much smaller amongst significant cis-SNPs because these are also concentrated near genes (right panel). (b) Each symbol corresponds to a gene-SNP association tested with observed or hidden genotypes respectively. For running BaseQTL with observed gentoypes, cis-SNPs are selected based on proximity to the gene under analysis as well as by having a minimum of 5 heterozygous individuals for the cis-SNP in the study sample. For running BaseQTL with hidden genotypes we selected proximal cis-SNPs with a minor allele frequency of at least 5% in the reference panel (1000 Genomes phase 3). As a result, in some occasions, for a given gene, the selection of cis-SNPs run with hidden genotypes and with observed genotypes may differ. To maximise the number of comparisons, we matched gene-SNP associations with r^2^≥ 0.9 between cis-SNPs run with different models, when possible. For simplicity only significant associations in at least one condition are shown, with the inset table summarising all associations tested. Dashed lines show 99% credible intervals. (c) Out of 73 genes tested, 10 had significant eQTLs with observed genotypes and 3 with hidden genotypes, all also significant with observed genotypes. (d) Example of a signal detected from 462 GEUVADIS individuals analysed by linear model^23^ captured with 86 samples and observed genotypes (upper panel) or hidden genotypes (lower panel) using BaseQTL. In each plot each symbol corresponds to a *MAPK1* cis-SNP within a 100KB window. The x-axis indicates the cis-SNP position and the y-axis corresponds to the −log_10_(p-value) reported for the 462 samples in GEUVADIS. Points are colored by significance. Associations not reported in the analysis of 462 individuals in the GEUVADIS study were considered not significant. To ease visualization, no significant associations in both datasets (“None”) are plotted with a p-value of 10^−4^ and those only called significant by BaseQTL with a p-value of 5×10^−5^. Using a linear model on the 86 samples tested by BaseQTL would have detected no significant results (minimum p=0.001).

In some occasions, for a given gene, the selection of cis-SNPs run with hidden genotypes and with observed genotypes may differ due to the different selection criteria in each method (**Figure 4b**). To maximise the number of comparisons, we matched gene-SNP associations with r^2^≥ 0.9 between cis-SNPs run with different models, when possible. Thus, we compared 1032 gene-SNP associations over 73 genes. The direction of the estimates for the eQTL effect was invariably consistent with that obtained when using observed genotypes (correlation 0.3, **Fig. 4b**). With hidden genotypes we detected 3 eGenes, out of the 10 with observed genotypes (**Fig. 4c**). We also checked whether the significant associations detected with hidden genotypes were also reported in our gold standard GEUVADIS dataset^23^. We found 80% of the significant gene-SNP associations (32/40) detected with hidden genotypes (2/3 eGenes) were also significant in the gold standard, though the imputation quality score had only limited influence in the positive predictive value (**Supplementary Fig.5**). However, for *NDUFA6* the same cis-SNP, rs55816780, is an eQTL in larger studies (>2000 individuals) in blood^26^, which may reflect gain of power from ASE. An example of an eQTL signal that was successfully captured with no previous knowledge of genotypes is shown in **Fig. 4d**.

### Novel skin eQTL in psoriatic and normal skin

Finally, we used our method to find eQTLs in a publicly available RNA-seq data set of 94 psoriasis skin samples and 82 controls^13^. To maximise discoveries relevant to psoriasis, we selected genes upregulated in psoriasis versus normal skin (51 genes^13^ and Online Methods), and/or within 100 kB of a psoriasis GWAS hit^27^ (380 genes). From the 429 unique selected genes, we were able to test ASE for 138, with 118 tested in both skin types, 16 in psoriasis only and 4 in normal skin only. After post-analysis QC which excluded putative hits for *SBSN* and *KRT6A* because the ASE signal was inconsistent across fSNPs (**Supplementary Figs. 9,11,12**, Online Methods), we found significant eQTLs for 21 genes: 8 in both conditions, 9 in psoriasis and 4 in normal skin only (**Fig. 5a**). Associations across 10 genes for the same SNPs were previously described in healthy skin^22^ or were previously reported eQTLs in psoriasis^28^ (**Fig. 5a and Supplementary Table 4)**, including *ERAP1*, *FUT2* and *RASIP1* which have eQTLs which are psoriasis GWAS-hits (rs30187 for *ERAP1*, rs492602 for *FUT2* and *RASIP1*)^27, 22^ (rs469758 and rs281379 proxies with r^2^=1 and 0.8, respectively).

**Figure 5.**
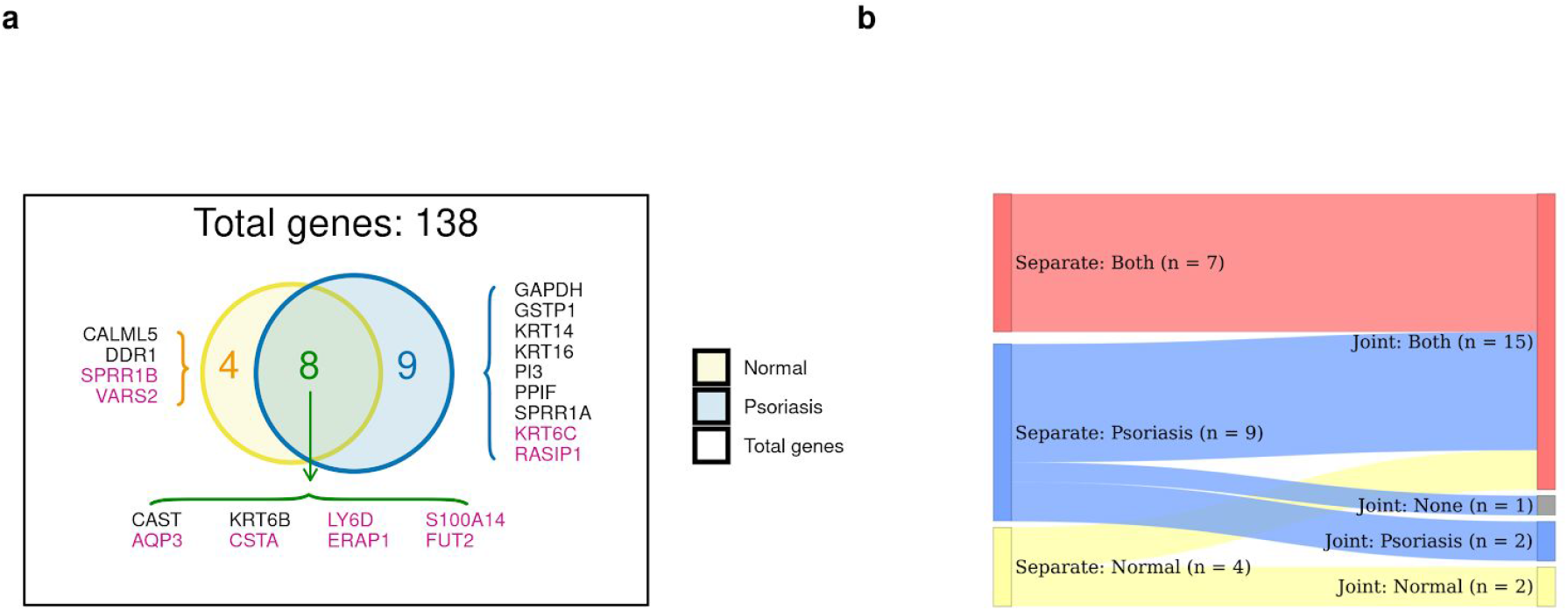
eQTLs in skin (a) Of 19 eGenes detected in analysis of either normal or psoriatic skin, only 8 were significant in both. For genes in pink, the same gene-SNP associations were also significant in GTEx for healthy skin (GTEx analysis V7). Moreover, associations for *ERAP1* and *FUT2* have been observed in a previous study of eQTLs in psoriasis^28^. (b) Sankey plot comparing results from running psoriasis and normal skin samples is separate models or jointly modelling eQTL effects. All the genes shown in (a) were run with the joint model except for *CSTA* which was excluded because only one cis-SNP was tested with normal skin and the significant signal observed in psoriasis could not be assessed in control samples (**Supplementary Table 5**). All 7 genes for which we observed signals in both tissues when run independently remained significant in both conditions with the joint model, and for eight of the 13 genes with apparent condition-specific effects, the joint model favoured a shared signal. Note that one psoriasis specific gene in the individual models (*SPRR1A*) was no longer significant in the joint model.

We exploited the flexibility available through using a standard statistical modelling language to jointly model eQTL effects in normal and psoriasis tissues in order to determine whether apparent psoriatic-specific effects reflected a lack of power in normal skin samples or were truly specific to psoriasis tissue (Online Methods). We restricted the analysis to 23 genes which were run in both skin types with significant associations at least in one (**Figure 5a**). Our joint model estimates two parameters: β_a_ which corresponds to the addition of the coefficients for the allelic fold-change in each skin and β_d_ which corresponds to their difference (Online Methods). We are particularly interested in β_d_ for assessing whether there is a difference in effects across conditions. Our prior for β_d_ expects half of eQTL signals shared in both tissues (1.5% weight) and half tissue specific (2 tissues, 1.5% weight on each), with 95.5% of null associations (Online Methods). All of the eGenes with common signals across skin types assessed with separate models were also shared with the joint model (Figure 5b). For the 9 specific psoriasis eGenes we detected with independent models, the joint model reported signals in both tissues for 6 of them, with *GSTP1* and *KRT14* specific for psoriatic skin, while eQTLs for *SPRR1A* were no longer significant (**Fig. 5b**). *PI3* is an antiproteinase and antimicrobial molecule highly upregulated in psoriasis ^13,29,30,13^ We detected an eQTL for *PI3* in psoriasis only when we ran separate models but the low expression in normal skin meant the joint model could not convincingly reject a common effect in both skin types (**Fig. 6a**). *GSTP1* is a representative example of a gene which appears specific for psoriasis skin. It is moderately upregulated in psoriasis (fold change = 1.7), so the psoriasis specific effect is unlikely to reflect lack of power on control samples (**Fig. 6b**). Of the 4 eGenes identified in normal skin only when running in separate models, the joint model confirmed specific signals in healthy skin for *SPRR1B* (**Fig. 6c**) and *DDR1* (**Fig. 5b**). The *SPRR1B* eQTL was also found in healthy skin samples from GTEx. Its strong upregulation in psoriasis (fold change 12) is therefore likely to be driven by an eQTL independent mechanism. Detailed plots for each gene can be found in **Supplementary Fig. 6–11** and summary results from the joint model can be found in **Supplementary Table 5.**

**Figure 6.**
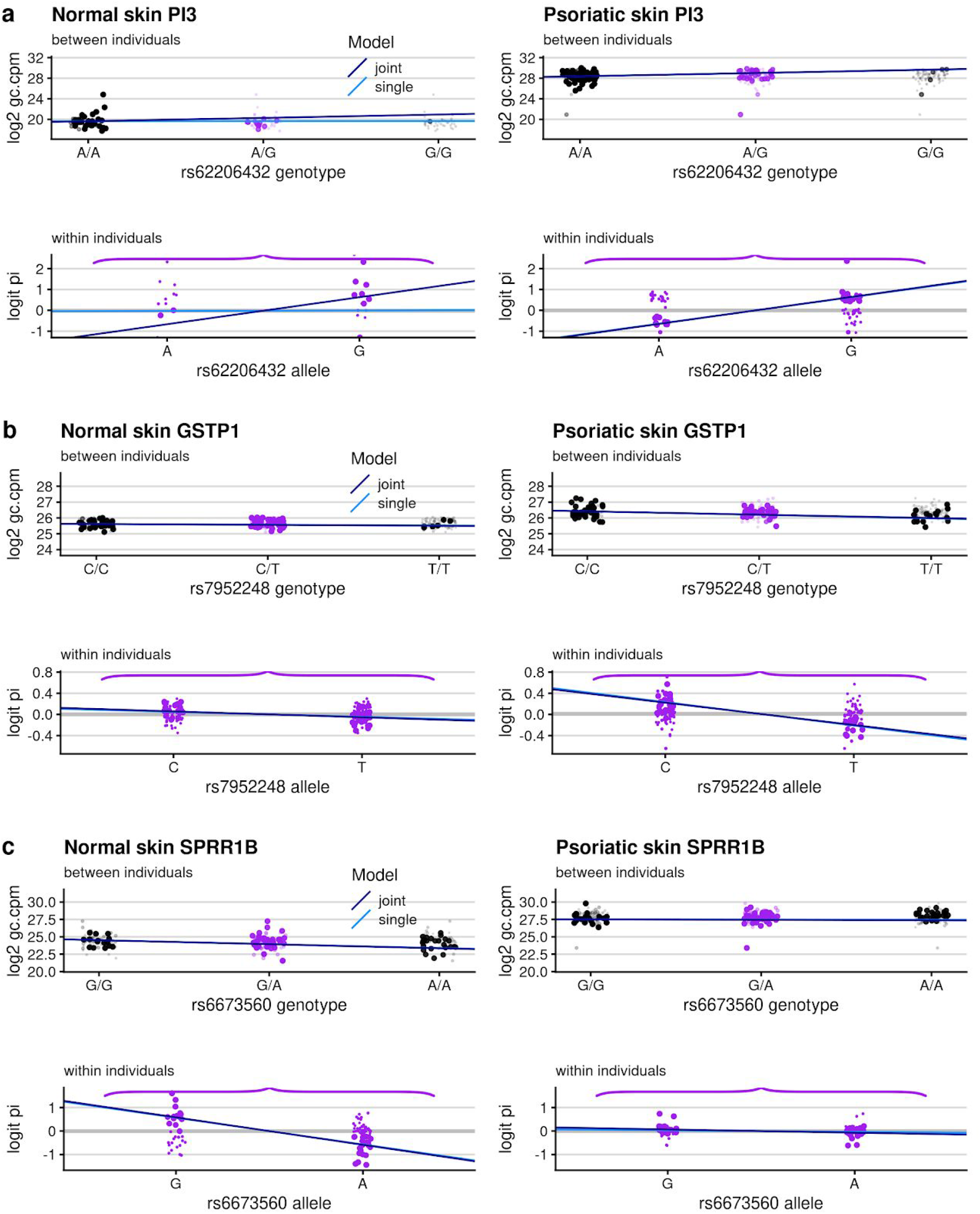
Disentangling condition specific eQTLs. eQTL estimates obtained from the joint or single models in normal or psoriasis skin (left and right sides). For each skin type we plotted the eQTL effect illustrating the two components of our likelihood: between and within individual variation (top and bottom panel, respectively). Between-individual plots show the genotype of the cis-SNP (x-axis) against the total gene counts per million reads, adjusted by GC-content (Online Methods) in log_2_ scale (y-axis). As genotypes are unobserved, for each individual we estimated the probability of each genotype and each point corresponds to the indicated genotype with the size and transparency indicating the probability. To represent within individual variation only heterozygotes are considered. The y-axis corresponds to the logit of the proportion of aggregated reads across fSNPs mapping the haplotype containing the alternative allele of the eQTL (as represented in **Fig.1**). The light and dark blue lines correspond to the mean effect obtained with the single or joint model, respectively. **a***PI3* **b***GSTP1* **c***SPRR1B*.

## Discussion

We have developed a novel statistical model, BaseQTL, for mapping genetic variants that regulate sequence-based molecular phenotypes. BaseQTL has increased power and positive predictive value over existing methods when tested for eQTL analysis and it is especially suitable for modest sample sizes.

Important differences between BaseQTL and alternative methods include the use of a reference panel for inferring haplotype phase and the use of a regularizing prior for shrinkage of effect estimates, to mitigate spurious extreme estimates which can arise from small sample sizes.^10,31^ In addition, contrary to WASP and RASQUAL which model ASE from haplotype pairs formed by the cis-SNP and each fSNP assuming independence, we aggregate haplotypic counts which may enable us to more faithfully model haplotype phase by accounting for the LD structure within genes. Last, we use a novel modelling strategy to account for reference mapping bias with the flexibility to model each fSNP independently without the need of simulations or discarding reads. All these factors may contribute to the improved performance of BaseQTL.

A major strength of BaseQTL is the ability to map eQTL in samples with no genotypes. Calling variants from molecular sequenced features combined with the use of an external reference panel for imputation allowed us to extract meaningful genetic signals from data sets of modest sample size. Although we expected the performance to be lower than with genotypes, when tested on a sub-sample of the GEUVADIS dataset we achieved 80% positive predictive value, assuming no false negatives in the full GEUVADIS analysis, with genotype imputation at the cis-SNP being the main source of false positives.

While BaseQTL provides strong flexibility with regard to cis-SNPs or fSNPs to run different types of sensitivity analysis and adapt to different analytical designs, it is computationally intense, with a median time of 6 and 10 minutes per gene for observed genotypes and hidden genotypes, respectively, using a cis-window of 1MB (**Supplementary Fig. 14)**. Thus, BaseQTL is particularly suited for targeted genomic regions to identify eQTL or condition specific eQTLs.

When we applied BaseQTL to skin we were able to validate associations for 10 out of 23 genes in normal skin from GTEx^7^ (Fig. 5a). Additionally, the signals we observed for *CAST,* and *GAPDH a*lthough not reported in normal skin, have been reported in blood ^26^, for *PPIF* in T cells ^32^ and for *KRT16* in adipose tissue (GTEx) (**Supplementary Fig.8**), composed of adipocytes, myeloid and lymphoid cells, among other cell types.

*GSTP1* is highly expressed in monocytes, dendritic cells and PBMC as a whole (https://www.proteinatlas.org/ENSG00000084207-GSTP1/tissue) and the signals we observed were specific to psoriasis and strongly significant in blood^33^ and neutrophils ^32^. Psoriatic skin is characterized by the proliferation of activated keratinocytes and infiltration of lymphocytes and myeloid cells ^34^. The fact that some of the eQTLs we observed were replicated in larger studies of blood but not in GTEx for normal skin could reflect gain of power from ASE modelling in our method, infiltration of immune cells driving psoriasis signals, or a combination of both. Further experiments using purified cell types from normal and psoriasis tissues will be instrumental to address these issues.

The flexibility offered by embedding our method within standard statistical software allowed us to disentangle condition specific effects. Overall, although our eQTL search was targeted psoriasis specific effects through its gene selection, joint modelling did not generally support condition specific effects identified by running psoriatic and normal skin samples separately, in agreement with recent studies showing substantial eQTL sharing among related cell types or tissues ^9,35^. This included putative psoriasis-specific effects from the separate models for *KRT6C* and *RASIP1*, which have reported signal in normal skin in GTEx. In addition, we found a novel eQTL for *PI3*, produced by epithelial and immune cells which regulates proliferation and inflammation ^36^. It is secreted by keratinocytes in response to IL-17 ^37^ or IL-beta and TNF-alpha ^38^, with protective functions against epidermis damage ^38^. PI3 levels in blood correlate with psoriasis severity ^39^.

BaseQTL can extract meaningful genetic signals from data sets of small sample size even without genotypes. We expect our method will facilitate discovery of cell type and disease-dependent eQTLs hidden in a wealth of RNA-seq data to unravel molecular mechanisms that contribute to disease.

## Acknowledgements

This work was supported by the Wellcome Trust (WT107881) and the MRC (MC_UU_00002/2, MC_UU_00002/4, MC_UU_00002/13, MR/R013926/1). S.W.R. was also supported by the NIHR Cambridge Biomedical Research Centre. The views expressed are those of the author(s) and not necessarily those of the NIHR or the Department of Health and Social Care.

## Author Contribution

C.W. conceived of the project. E.V. C.W. and S.W.R. developed the model. E.V. wrote the software and performed analyses. W.Y.L. C.S. performed analyses and implemented the software. P.D.W.K. and S.W.R. contributed to the design of statistical analysis. E.V. and C.W. wrote the manuscript with input from all authors. C.W. directed the project.

## Online Methods

### Code and URLs

The source code and documentation for BaseQTL are open source available at https://gitlab.com/evigorito/baseqtl_pipel which includes a pipeline to process RNA fastq files and genotypes, if available, to prepare for running BaseQTL (Supplementary Fig.13).

GEUVADIS samples were accessed from E-GEUV-1, ftp://ftp.sra.ebi.ac.uk/vol1/fastq Psoriasis and normal skin samples were accesed from E-GEOD-54456, ftp://ftp.sra.ebi.ac.uk/vol1/fastq

GTEx associations for skin, blood and lymphoblastic cell lines corresponding to Analysis V7 were downloaded from https://gtexportal.org/home/datasets.

Deferentially regulated genes between psoriasis and normal skin were downloaded from https://ars.els-cdn.com/content/image/1-s2.0-S0022202X15368834-mmc2.xls

### Data samples

We downloaded RNA-seq data from 86 GEUVADIS samples with EUR ancestry (GBR code) from ArrayExpress (E-GEUV-1, Supplementary Table 6). We also analysed 94 and 90 RNA-seq normal and psoriasis skin samples [1] obtained from ArrayExpress (E-GEOD-54456, downloaded 2/11/2018). For the analysis of psoriasis eQTL we selected up-regulated in psoriasis versus normal skin (51 genes based p≤10-6 (corresponding to family-wise error rate < 0.025) and a median expression of at least 500 RPKM in psoriasis samples (data extracted from https://ars.els-cdn.com/content/image/1-s2.0-S0022202X15368834-mmc2.xls,[1]), and/or within 100 kB of a psoriasis GWAS hit [2] (380 genes).

### RNA-seq preprocessing

For the psoriasis and normal skin RNA-seq data quality control using FASTQC indicated a high number of reads with Ns, which were filtered out using Prinseq [3]. All samples were aligned to the human genome assembly GRCh37 using STAR [4], but 3 of the normal samples failed alingnment and were excluded from downstream analysis. We calculated gene expression abundance by overlapping reads to an union of annotated Ensembl exons, excluding reads overlapping different genes as we did not have strand information.

### Calling genotypes from RNA-seq and phasing

For calling SNPs we fed the aligned files into bcftools [5] selecting uniquely mapping reads with a quality score of at least 20. We kept variants with read depth at least 10 that were also reported in the 1000 Genome project version 3 (haplotypes from 2504 individuals in NCBI build 37 (hg19) coordinates from mathgen.stats.ox.ac.uk/impute/1000GP_Phase3.html downloaded on 26/1/2018), with minor allelic frequency at least 0.05 in European individuals.

We assessed genotype errors in the fSNPs by comparing the genotype frequency of each fSNP in the samples relative to samples of same ethinicity in the reference panel by Fisher’s exact test. We report the p-value for each fSNP. Moreover, we discard SNPs with minor allele frequency below and we set a cut-off of at least 100 reads before second alignment, but those are user defined thresholds.

### Quantifying the number of reads overlapping fSNPs

We adapted phASER [6] to count the number of reads overlapping each fSNP. We followed the guidelines for ASE quantification suggested by Castel et al (genome biology 2015 16:195), by restricting the analysis to uniquely mapped reads with base quality for fSNPs ≥ 10. We first used the ‘phaser.py’ command that count reads overlapping SNPs. Phaser requires phased genotypes as input, so we used SHAPEIT2 [7] using the 1000 Genomes phase3 reference panel of haplotypes. Next, we adapted phASER function ‘phaser_gene_ae.py’ to count only once reads overlapping two or more heterozygous variants.

We implemented the following QC steps to minimise false calls: first, when no strand information is available from RNA-seq we only considered fSNPs uniquely mapped to one gene. Second, we only use reads uniquely mapped reads to a locus and we correct for double counting of reads overlapping more than one fSNP.

### Statistical model

Our model maps QTLs for genetic variants (cis-SNPs) within a chosen distance to a feature (gene, isoform, ChIP-peak). For each feature, we consider all SNPs within it together with one potential regulatory SNP (cis-SNP), we jointly modelled the total read counts in the feature and the allelic imbalance between the chromosomes carrying the cis-SNP and the fSNPs. Our model builds up from the TrecASE model [8] by allowing phasing uncertainty, modelling reference panel bias and unobserved genotypes in a Bayesian framework.

### Basic model: known phase and genotypes

We begin by summarising our implementation of the TRecASE model [8] with observed genotypes and fixed phasing (Fig. 1a). The likelihood can be decomposed into a product of contributions from between individual (*L*_between_) and within individual (*L*_within_) likelihoods.

Let *c*_*i*_ be the total read counts at the specific feature for individual *i* (*i* =1,…,N), *G*_*i*_ the number of alternative alleles at the cis-SNP (0,1,2) and **x_i_** a vector of *p* covariates. We used the same parametrization as in TRecASE. We modelled total gene counts *c*_*i*_ by a negative binomial distribution (*f*_*NB*_) to allow for over-dispersion of RNA-seq reads:

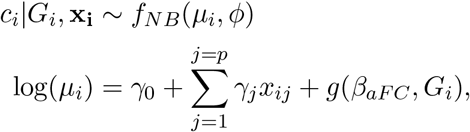

**Where do we say what covariates are used in the real data analysis?** where *g*(*β*_*aFC*_, *G*_*i*_) models the genetic effect and *β*_*aFC*_ corresponds to the expected log-allelic fold change of individuals homozygous for the alternative allele to those homozygous for the reference allele for the tested cis-SNP, as defined by the TReCASE model (ref):

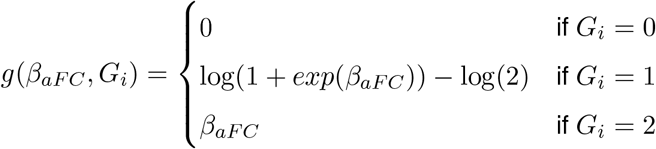

To model allele-specific expression (ASE) we assume initially that we observe for each individual *i* their complete haplotypes formed by the cis-SNP and fSNPs. We distinguish these as (*h*_0*i*_, *h*_1*i*_) according to the haplotype carrying the reference and alternative alleles at the cis-SNP, respectively for individuals heterozygous at the cis-SNP, or arbitrarily for homozygous individuals. We count reads mapping to each haplotype by aggregating the counts across heterozygous fSNPs, according to their phase, in each individual. Of *m*_*i*_ ≤ *c*_*i*_ reads which overlap at least one heterozygous fSNP, we denote by *n*_1*i*_ the number which map to *h*_1*i*_. We model *n*_*i*1_|*m*_*i*_ by a beta-binomial distribution with *π* being the expected proportion of ASE and *θ* the overdispersion parameter as follows:

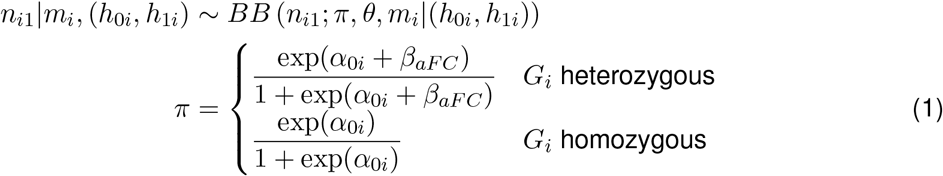

where *α*_0*i*_ is a random intercept parameter which depends on (*h*_0*i*_, *h*_1*i*_) (although we drop the dependence in the notation above for simplicity) and which would be 0 in the absence of reference sequence mapping bias. Homozygous individuals for the cis-SNP carrying heterozygous fSNPs contribute information for estimating the overdispersion parameter *θ*. The within-individual likelihood can therefore be expressed as:

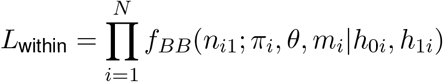

We put uninformative priors on the standard regression parameters

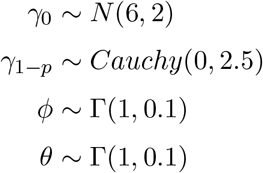

and describe informative priors for *α*_0*i*_ and *β*_*aFC*_ in sections below.

### Extension 1: Modelling phasing uncertainty

We now relax the assumption that (*h*_0*i*_, *h*_1*i*_) are observed and known. We denote by *F*_*i*_ the unphased genotypes across the fSNPs for individual *i*, and by *G*_*i*_ their genotype at the cis-SNP. We account for phasing uncertainty by averaging over the likelihood of *n*_1*i*_|(*h*_0*i*_, *h*_1*i*_) over all possible 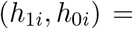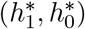, weighted by a simple maximum likelihood estimate of 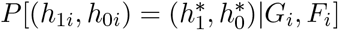 estimated from 1000G reference panel phase 3. Thus the likelihood contribution becomes

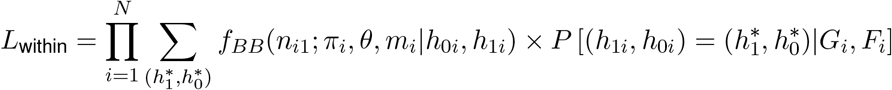

### Extension 2: Unknown cis-SNP genotypes

We use the same idea to consider *G*_*i*_ latent, deriving the haplotype pair probabilities conditional only on the observed *F*_*i*_, since *G*_*i*_ is directly specified by any haplotype pair.

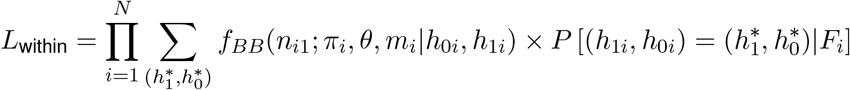

but we also need to adjust

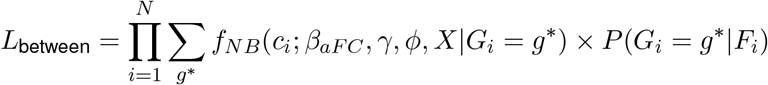

We use a standard measure of imputation quality (*r*^2^) [9]. If *G*_*i*_ ∈ {0, 1, 2} and *p*_*ik*_ = *P* (*G*_*i*_ = *k*|*F*_*i*_) is the probability obtained by imputation that the genotype of the *i*th individual is *k* (section Extension 2: Unknown cis-SNP genotypes), the expected allele dosage for individual *i* is *E*(*G*_*i*_) = *p*_*ik*=1_ + 2*p*_*ik*=2_. The information metric is defined as

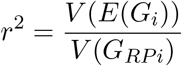

with *G*_*RPi*_ the genotype for the cis-SNP for individual *i* in the reference panel. We report this value in the summary output.

### Extension 3: jointly modelling different conditions in unpaired samples

We describe this in context of our application to psoriatic and normal skin, but the same method applies to compare unpaired data from any two conditions or cell types. We can write the between individual component of independent models for normal (N) and psoriasis (P) skin as:

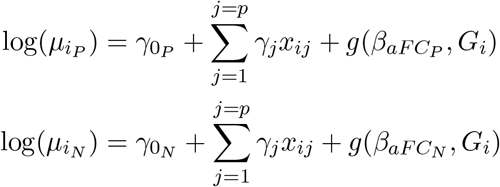

We can jointly model total gene counts from normal and psoriasis skin as follows:

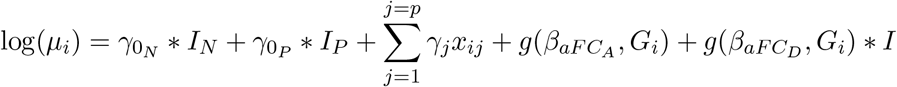

With:

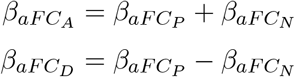

And:

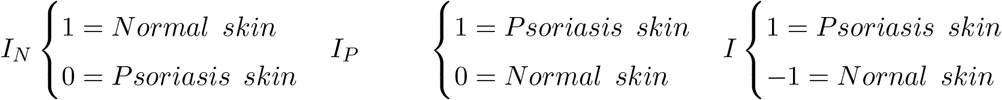

Rather than the more usual treatment contrasts, using zero/one dummy variables, we use sum-to-zero contrasts for the group variable. Mathematically the models are identical, there is only a change in interpretation of the resulting coefficients. As a difference in conditions is our primary focus, the sum-to-zero contrast will directly assess whether there is any condition difference (regardless of direction/sign and interactions).

### Prior specifications

#### Modelling reference mapping bias, prior on *α*_0*i*_

To estimate expected reference panel bias at each fSNP *k*, we pooled observed and pseudo reads across all individuals. Let *r*_*k*_ and *t*_*k*_ be the number of reads re-aligned to the alternative allele and the total number of re-aligned reads, respectively. Thus, 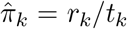 is the proportion of reads mapping to the alternative allele. On rare occasions we observed 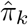 higher than 0.5. Often when this happened, two or more SNPs were close to each other and shared overlapping reads with some alleles being reference and other alternative in the original read. We apply a binomial test assessing whether the bias estimate is significantly higher than 0.5 and discard those fSNPs with *p* < 0.01 because this pattern was not observed in the distribution of bias estimated from the observed reads only.

When *β*_*aFC*_ = 0, then logit(*π*) = *α*_0*i*_ (1). Note that the effect of any bias in our likelihood will depend on how the alternative allele at the fSNP is phased with the alternative allele at the cis-SNP. Let 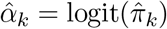, and define

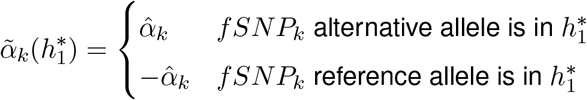

We assume that log(*α*_0*i*_|(*h*_0*i*_, *h*_1*i*_)) is normally distributed, with expected value a weighted average of 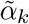

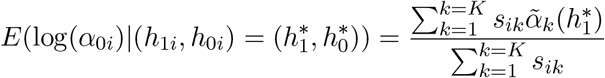

where *s*_*ik*_ is the number of reads in sample *i* overlapping fSNP *k*, and variance

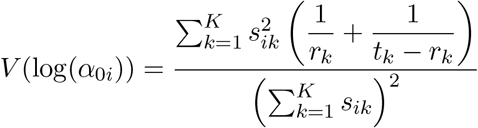

This is then an informative prior for *α*_0*i*_ that captures both local sequence effects and variable coverage between individuals and between SNPs, derived from observed data and its counterfactual alternative.

#### Informative prior on *β*_*aFC*_

We also use a data-derived prior on *β*_*aFC*_, building on information amassed from large eQTL studies. We used estimates of eQTL effects from cis-SNPs in GTEx, assuming true eQTL effects, 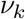 at each SNP *k*, come from a mix of Gaussian distributions

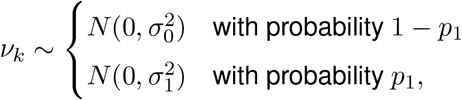

where 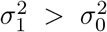, and that estimated eQTL effects are unbiased, i.e.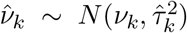 where 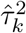 is the standard error of the eQTL effect such that for SNP/gene pair *k*, 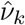 We took a sample of 10^6^ 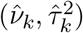 values for unlinked SNPs within 1 Mb of the target gene’s transcription start site. We estimated 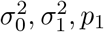 by Metropolis-Hastings, and code to run this analysis is available at https://github.com/chr1swallace/fitmix.

#### Informative priors on *β*_*aFC*_*A*__ and *β*_*aFC*_*D*__

We can express the prior for *β*_*aFC*_ as:

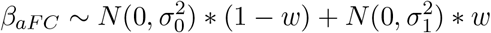

When jointly modelling normal (N) and psoriasis (P) skin we have:

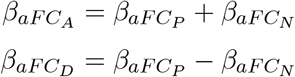

Under independence:

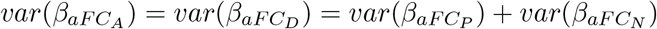

The priors for *β*_*aFC*_*A*__ and *β*_*aFC*_*D*__ can be expressed as a mixture of Normal distributions with the following components:

| eQTL effect | Mean | Variance | Weight |
| --- | --- | --- | --- |
| Neither | 0 | $2\sigma_0^2$ | 95.5 |
| Either | 0 | $\sigma_0^2 + \sigma_1^2$ | 3 |
| Both | 0 | $2\sigma_1^2$ | 1.5 |

### Running linear model RASQUAL and BaseQTL

We ran the linear model in R using “lm” function regressing the logarithm of total gene counts on genotypes.

For RASQUAL we created a VCF file with allele specific counts using the tools provided in https://github.com/kaurala We run RASQUAL with normal settings or the permutating test.

BaseQTL inputs were prepared as detailed above and at https://gitlab.com/evigorito/baseqtl_pipeline. All three models were adjusted by GC-corrected library size as implemented in the library rasqual-Tools.

### Calculation of FDR

For the linear model we calculated FDR using the R function “p.adjust” with method “BH”. For RASQUAL we used the method provided by RASQUAL itself [10], namely to generate a single p value from muted data, and defining

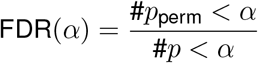

where *p*_perm_ corresponds to the permutation p values, *p* to observed data p values, and *α* the significance threshold.

For BaseQTL, we calculated 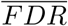 as described in [11]. Briefly, they define FDR as:

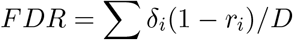

where *δ*_*i*_ is an indicator for rejecting the i-th eQTL comparison, *D* = ∑*δ*_*i*_ corresponds to the number of rejections and *r*_*i*_ ∈ {0, 1} denotes the unknown truth for a SNP being (1) or not (0) a cis-eQTL. While *r*_*i*_ is unknown, we can calculate *v*_*i*_ = *P* (*r*_*i*_ = 1|data) from our posterior samples by calculating the proportion of times that 0 was excluded from credible intervals of specific size (*α*′ = 85%, 90%, 95% and 99%). (To do this with a manageable number of samples, we used normal approximations to the marginal posterior distribution of the eQTL effect). Under those conditions using a cis-window of 1MB or 0.1MB we observed the following 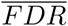:

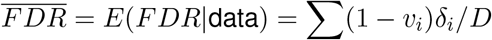

| Decision rule | D (1MB) | D (0.1MB) | $\overline{FDR}$ |
| --- | --- | --- | --- |
| null $\notin$ 99% CI | 193 | 153 | 0.001 |
| null $\notin$ 95% CI | 347 | 262 | 0.013 |
| null $\notin$ 90% CI | 846 | 511 | 0.051 |
| null $\notin$ 85% CI | 2084 | 977 | 0.097 |

### Definition of significant associations using BaseQTL

We defined significant associations those for which 0 was excluded from the 99% credible intervals of the posterior distribution, unless otherwise stated. This threshold was a good compromise between positive predictive value and sensitivity (Fig.3).

### Consistency of the ASE signal across fSNPs

We implemented a quality control measure aiming to exclude potential false positive calls. Given f fSNPs within a gene, for each fSNP we extracted the number of reads mapping the alternative allele in each individual (*n*_*i*_) and the total number of reads mapping the fSNP (*m*_*i*_). We then fit the following models:

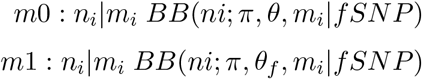

We use an anova test to compare the models and set a p-value threshold of 0.05 to exclude genes from analysis if there is evidence of difference in the over-dispersion parameters across fSNPs. We run the analysis using the “betabin” function from the “aod” R library.

## Supplementary note

### Mitigating genotype errors

To minimize genotype errors performed the following steps: First, we called variants using a base quality threshold above 20 and limiting the analysis to uniquely mapped reads. Second, we filtered out called variants not annotated in the external reference panel (1000 Genomes phase3). Third, by excluding variants according to a depth threshold. Ideally, we would be calling genotypes for the same fSNPs that were used for inference with observed genotypes, which were 498 fSNPs across 86 samples, adding to 42,828 calls calls. We have chosen to limit RNA-seq calls to those fSNPs with depth ≥10, as this value provided a good trade-off between error rate and missing calls (Supplementary Fig. 2a). Last, by excluding fSNPs with different rates of heterozygosity across study samples compared to European samples from 1000 Genomes when performing a Fisher exact test (Online Methods). We selected a p value of 0.01 as a good compromise to exclude fSNPs with a high proportion of errors (Supplementary Fig. 2b). Using this threshold we excluded 2 fSNPs with 33 errors out of 82 with 4 missing calls and 8 out of 20 with 66 missing calls. Last, we looked at whether fSNPs with a higher number of missing calls across samples were associated with higher error rates. This was not the case (Supplementary Fig. 3a), so we did not exclude fSNPs based on the number of missing calls.

---

**Supplementary Figure 1.**
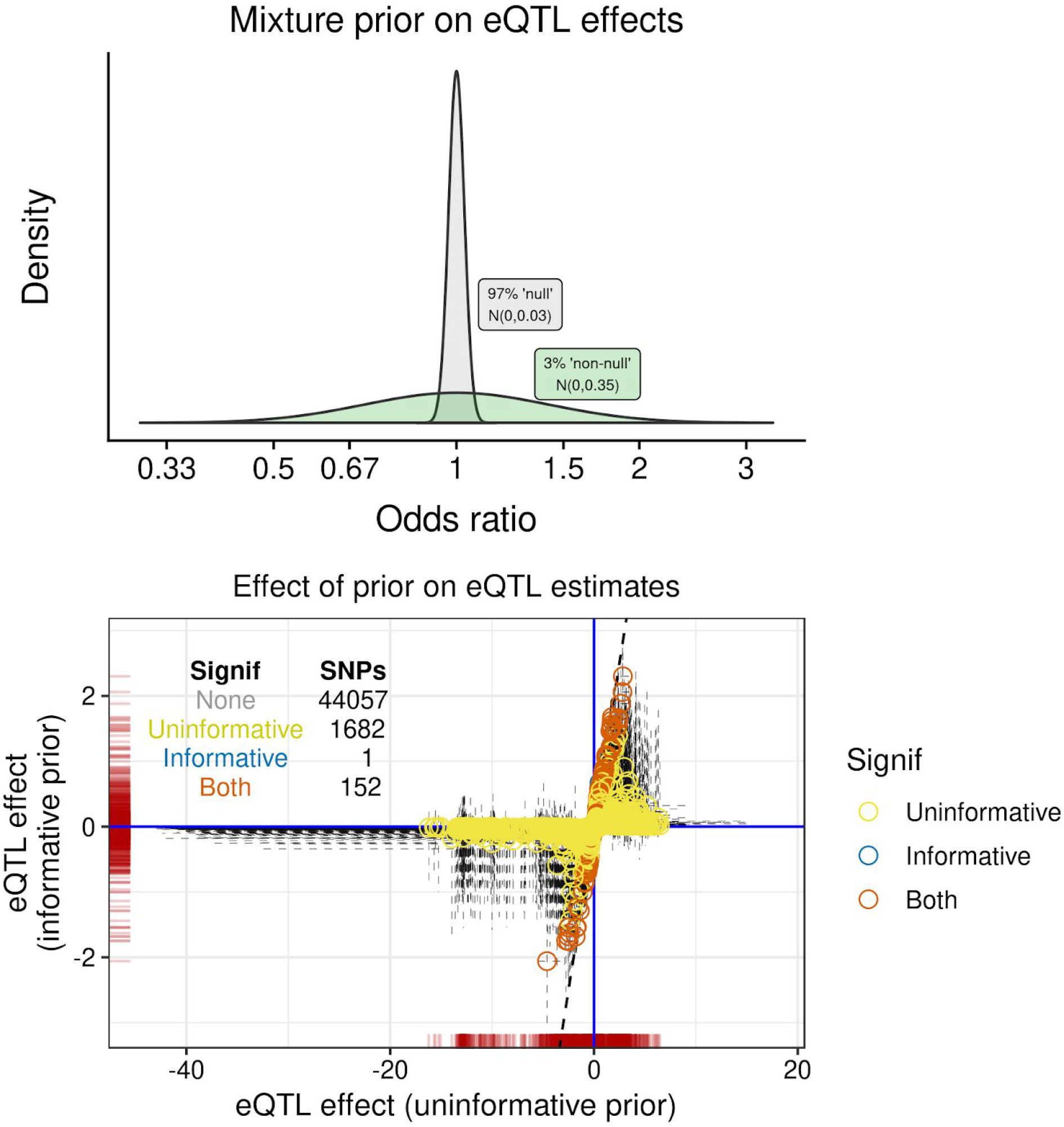
Shrinkage effect of prior on eQTL estimates. (a) we learnt an informative prior on eQTL effect sizes from GTEx LCL which is a mixture of a narrow (97%) and a wider (3%) central normal distributions, with sd=0.03 and 0.35 respectively.(Online Methods). (b) BaseQTL was run twice, once with this informative prior and once with an uninformative prior (N(0, 100)). The informative prior shrinks 92% (1682/1834) of significant effects so they are no longer significant, which changes the positive predictive from 0.25 to 0.90 when using the larger GEUVADIS dataset of 462 individuals as gold standard. Each point point corresponds to the eQTL effect (log_2_ allelic fold-change) running BaseQTL with observed genotypes for expressed genes in chromosome 22 (264) with the informative prior we derived (y-axis) or an uninformative (x-axis) prior. The gray lines indicate 99% credible intervals.

**Supplementary Figure 2.**
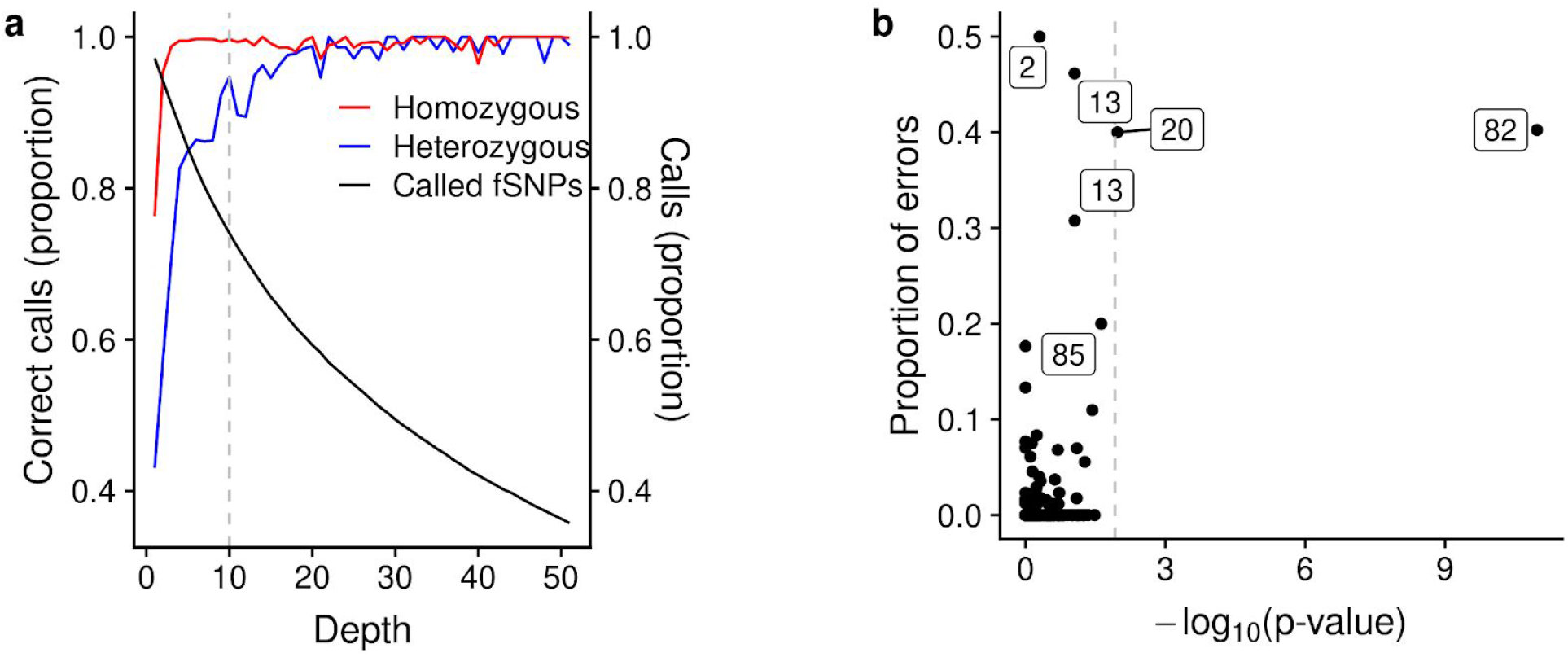
Quality control of RNA-seq genotyping errors. **(a)** Trade-off between genotype accuracy and number of variants called. Genotype concordance for calling fSNPs with RNA-seq or short read DNA genotyping increases with read depth for homozygous or heterozygous SNPs (red and blue lines with left y axis), while the proportion of variants with genotype calls decreases (black line with y right axis). **(b)** Each symbol corresponds to a fSNP genotyped across the 86 samples. The x-axis shows the −log10 p-value obtained by comparison of the frequency of heterozygous individuals relative to a reference panel of the same ethnicity (Online methods). The y-axis indicates the proportion of genotyping errors across samples when calling genotypes with RNA-seq relative to DNA sequencing. The labels indicate the total number of samples with genotypes called by RNA-seq for the fSNPs with the highest proportion of errors. The dashed vertical line at x-axis=2 (p-value = 0.01) is the threshold we selected.

**Supplementary Figure 3.**
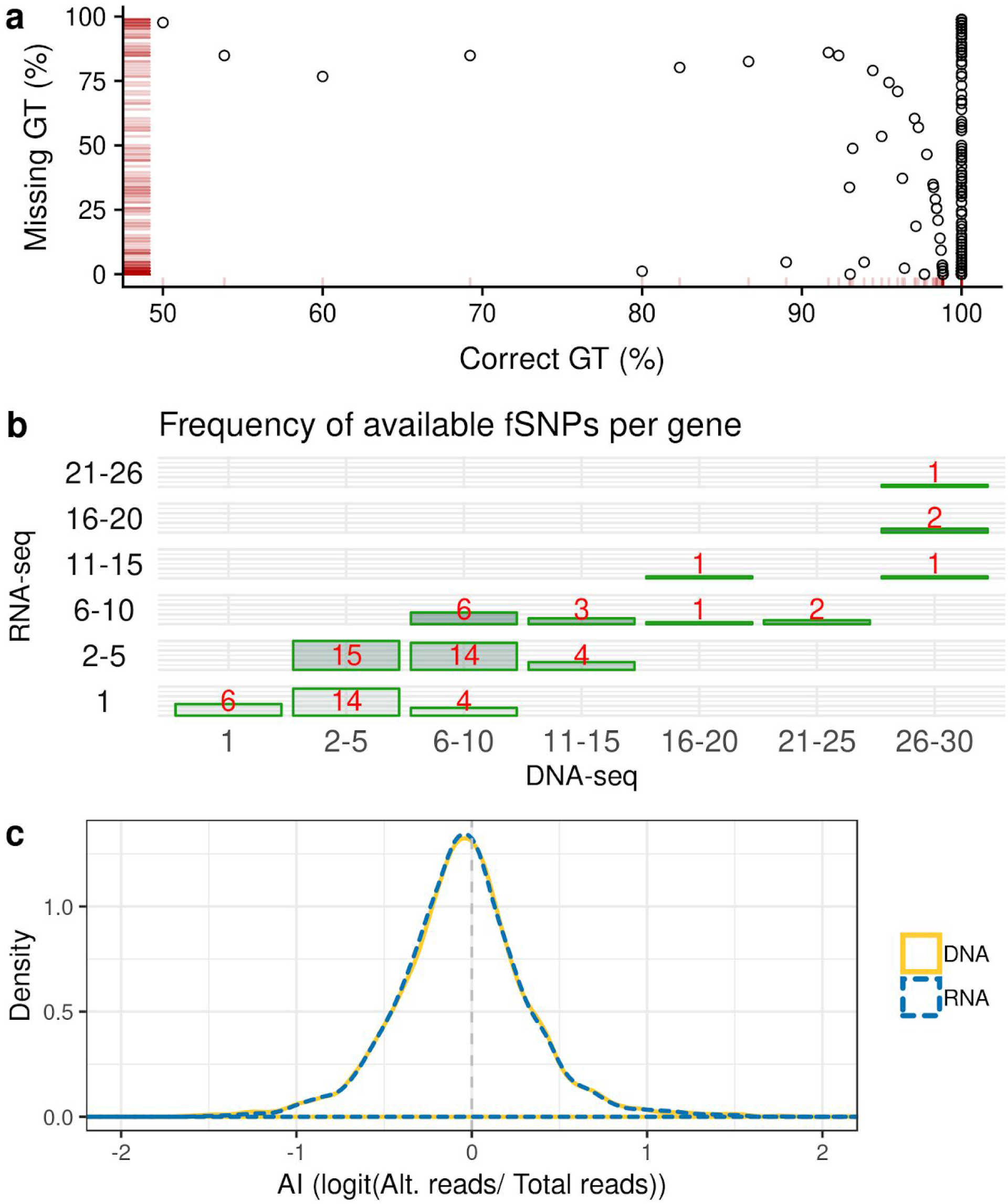
Genotyping fSNPs by RNA-seq. (a) Each symbol corresponds to a fSNP. The plot shows the proportion of samples with same genotype calls in DNA-seq and RNA-seq (x-axis) relative to the proportion of individuals with missing genotypes in RNA-seq calls (y-axis). As genotype errors were independent of missing values, we did not apply a threshold based on the number of missing genotypes (b) RNA-seq genotyping reduces the number of available fSNPs per gene. For each gene, the number of fSNPs used for inference was categorized as 1, 2-5, 6-10, 11-15, 16-20, 21-25, 26-30 both for observed or hidden genotypes. The bars correspond to the number of genes for a given number of fSNPs. (c) Distribution of raw AI estimates at each fSNP are similar between DNA-seq or RNA-seq. The dashed line at logit AI=0 corresponds to no imbalance.

**Supplementary Figure 4.**
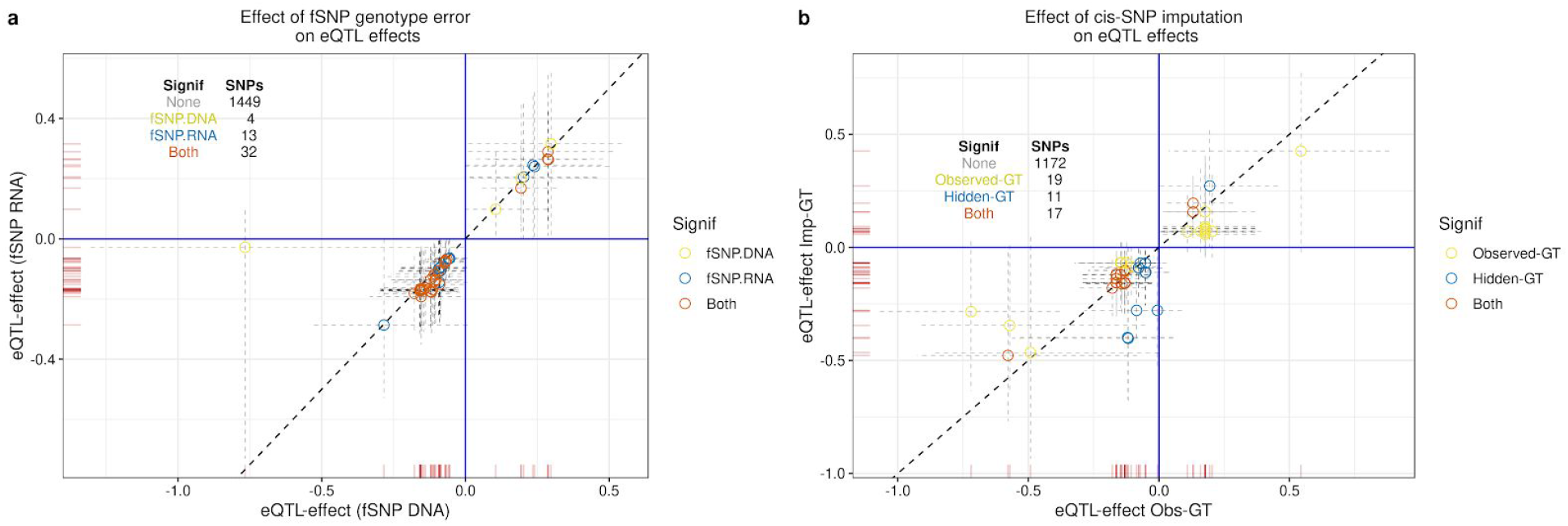
Dissecting the effect of genotyping errors on eQTL estimates. When running BaseQTL with hidden genotypes for the cis-SNP we restricted the analysis to cis-SNPs with a quality of imputation ≥ 0.3. (a) BaseQTL was run with fSNPs genotyped by DNA-sequencing or by RNA-seq. In both cases the same fSNPs were used for inference and the cis-SNP was imputed. Each symbol corresponds to the eQTL effect (log_2_) comparing both conditions with the dashed lines indicating the 99% credible intervals. (b) Same as (a) except that BaseQTL was run with fSNPs genotyped by DNA-sequencing and the genotype for the cis-SNP was either observed or imputed.

**Supplementary Figure 5.**
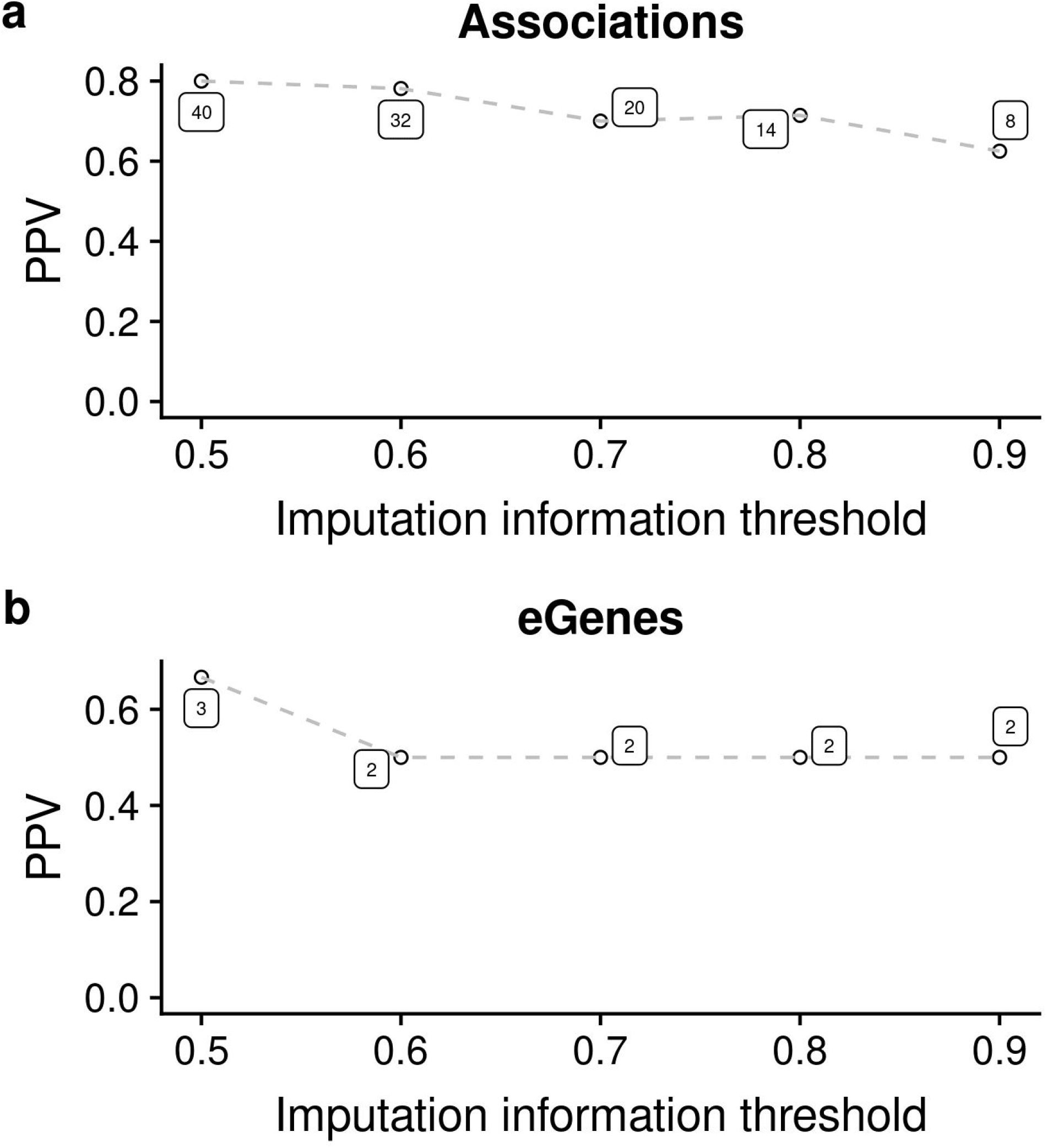
Comparing associations detected with hidden genotypes on a sub-sample of 86 individuals relative to a large GEUVADIS study of 462 samples. At each threshold of imputation quality (x axis) the PPV for associations (a) or eGenes (b) is shown. The values on the graph correspond to the total number of associations (a) or eGenes (b) called significant with hidden genotypes.

**Supplementary Figure 6.**
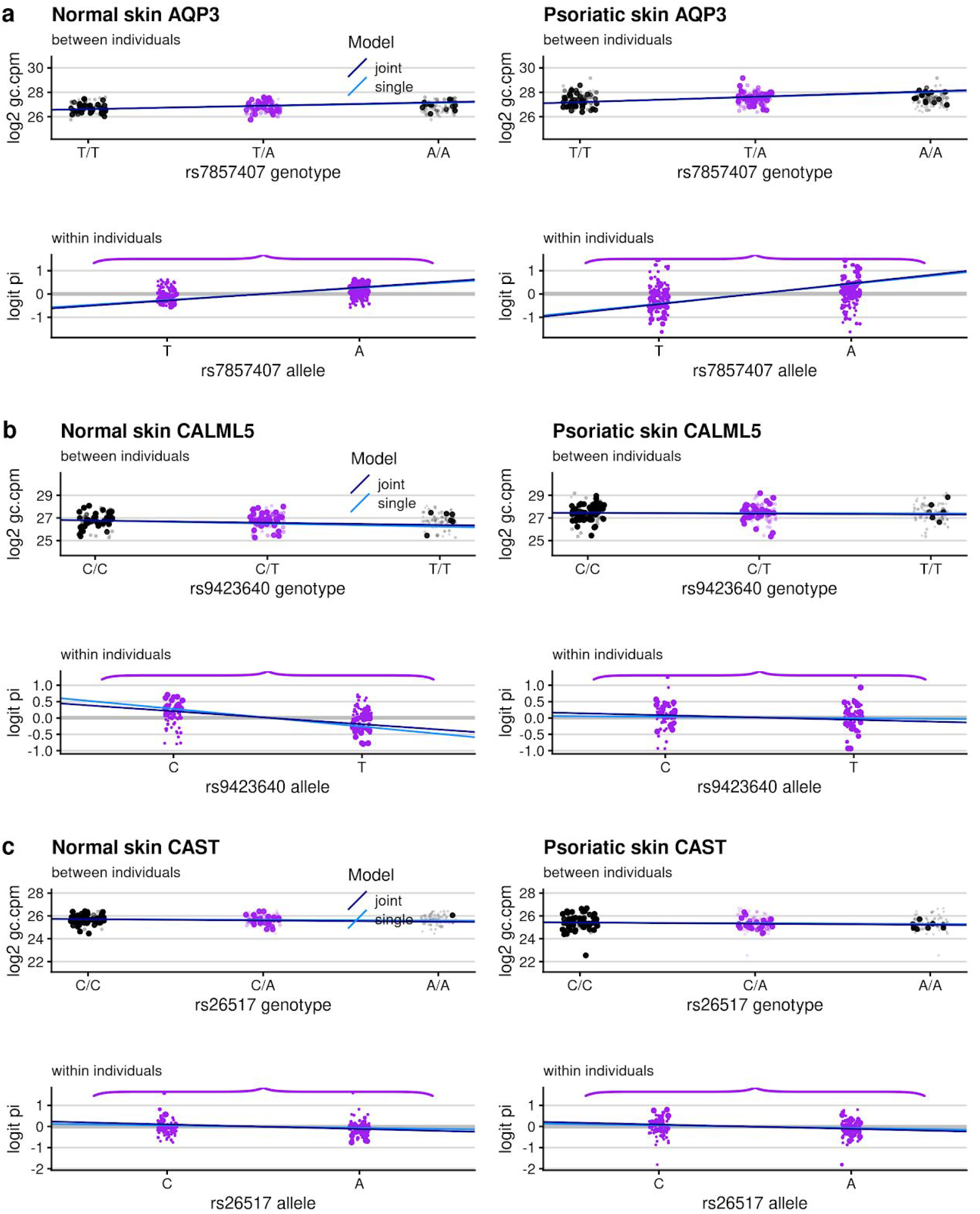
eQTL examples for the indicated genes. Same analysis as in Figure 6.

**Supplementary Figure 7.**
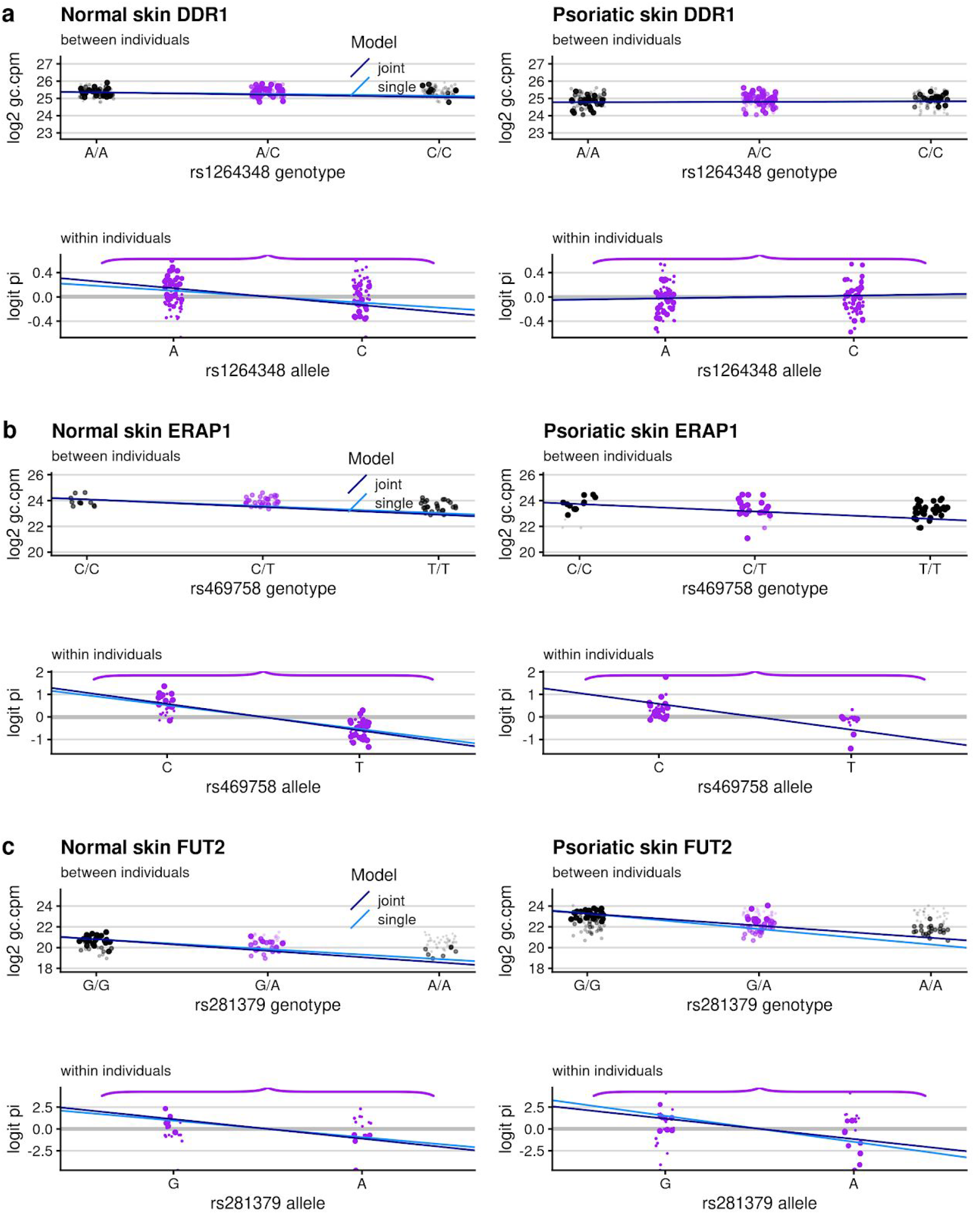
eQTL examples for the indicated genes. Same analysis as in Figure 6.

**Supplementary Figure 8.**
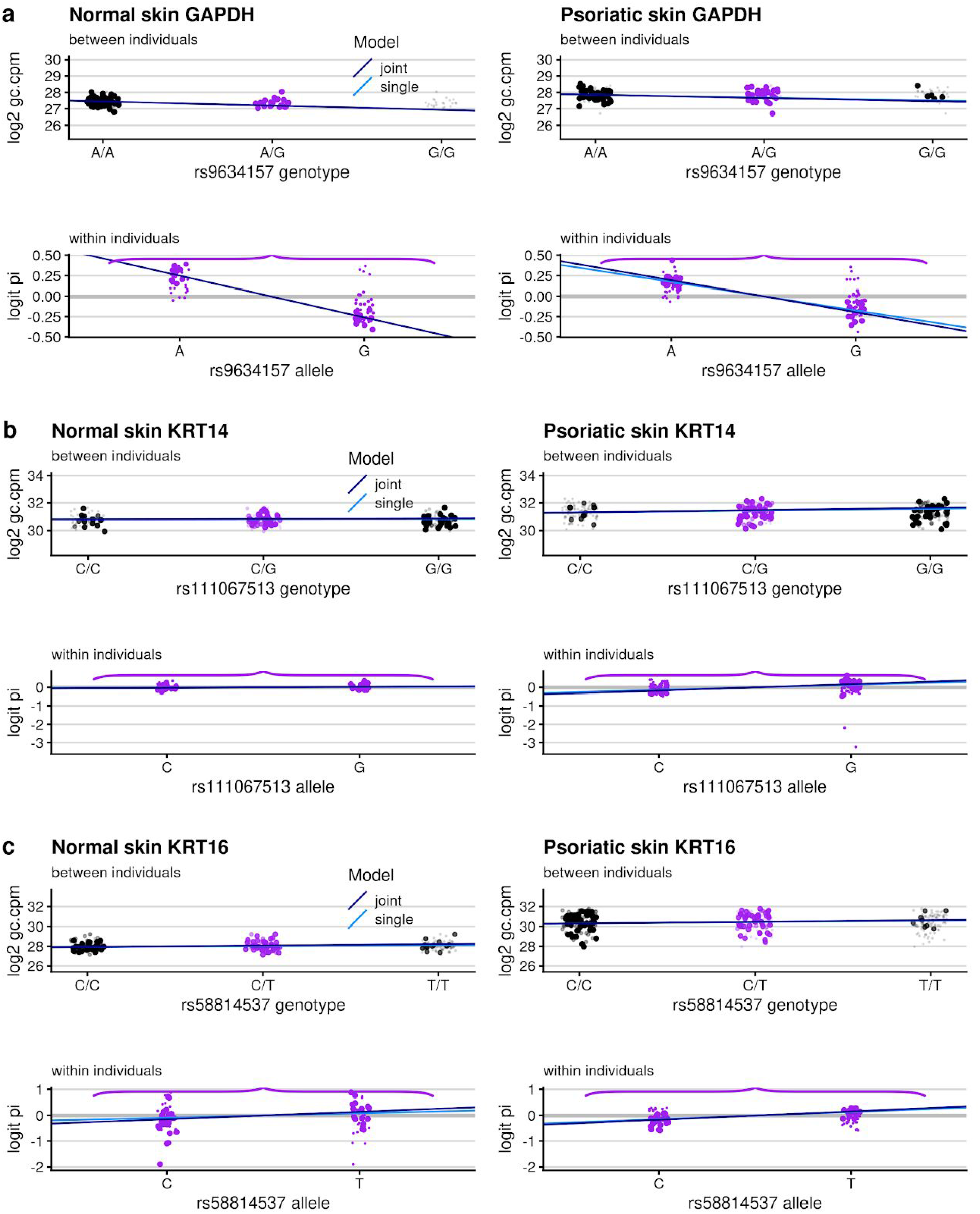
eQTL examples for the indicated genes. Same analysis as in Figure 6.

**Supplementary Figure 9.**
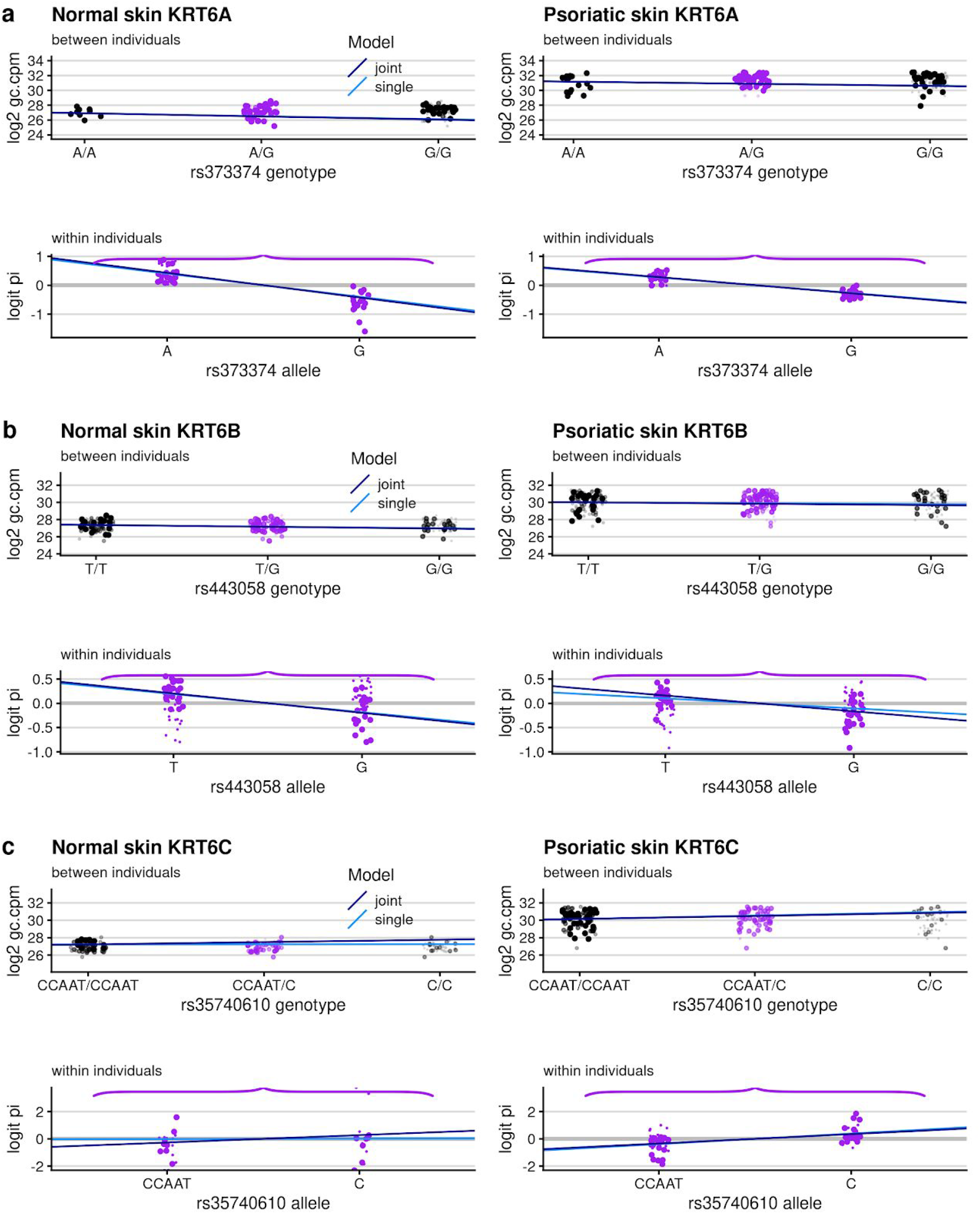
eQTL examples for the indicated genes. Same analysis as in Figure 6.

**Supplementary Figure 10.**
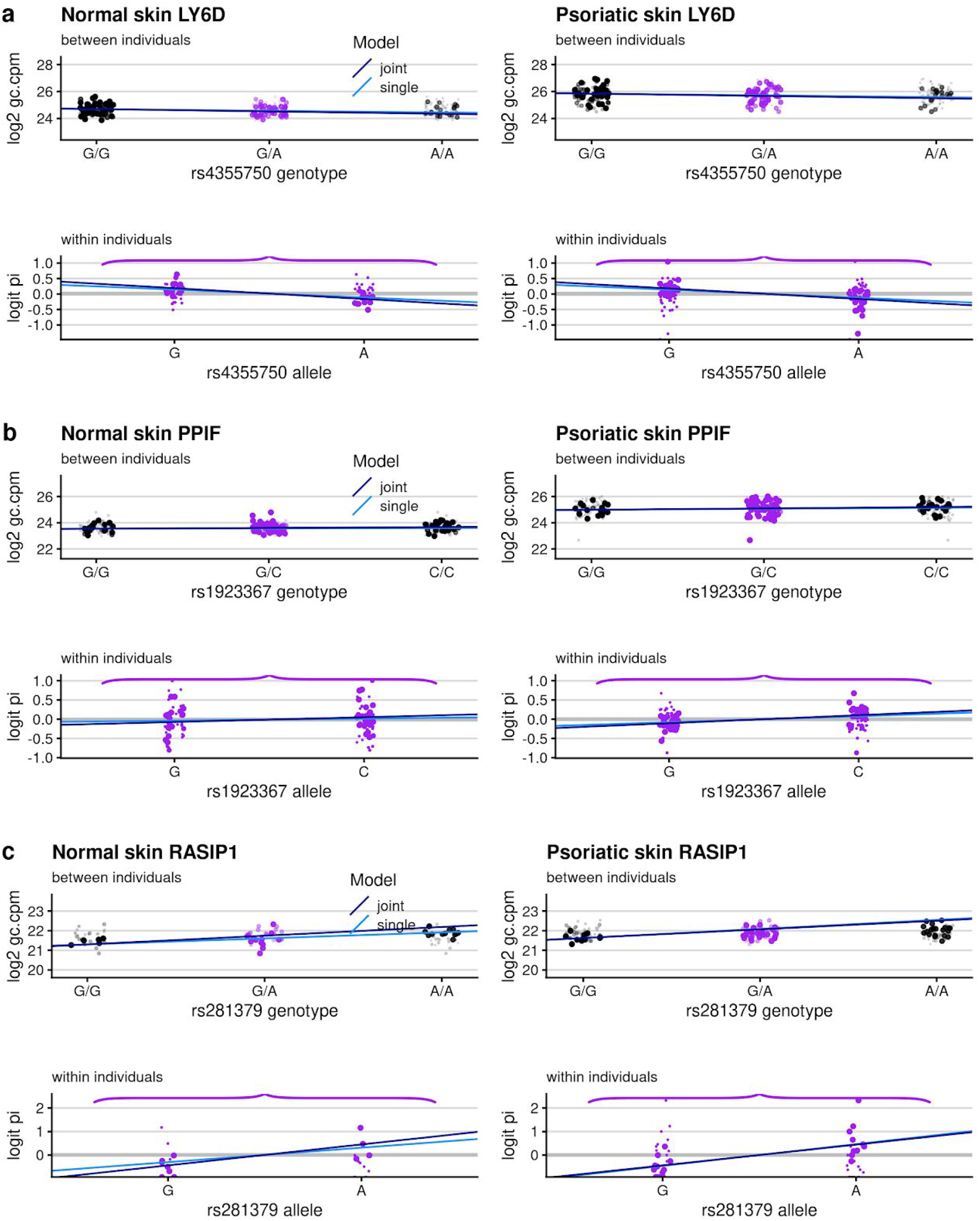
eQTL examples for the indicated genes. Same analysis as in Figure 6.

**Supplementary Figure 11.**
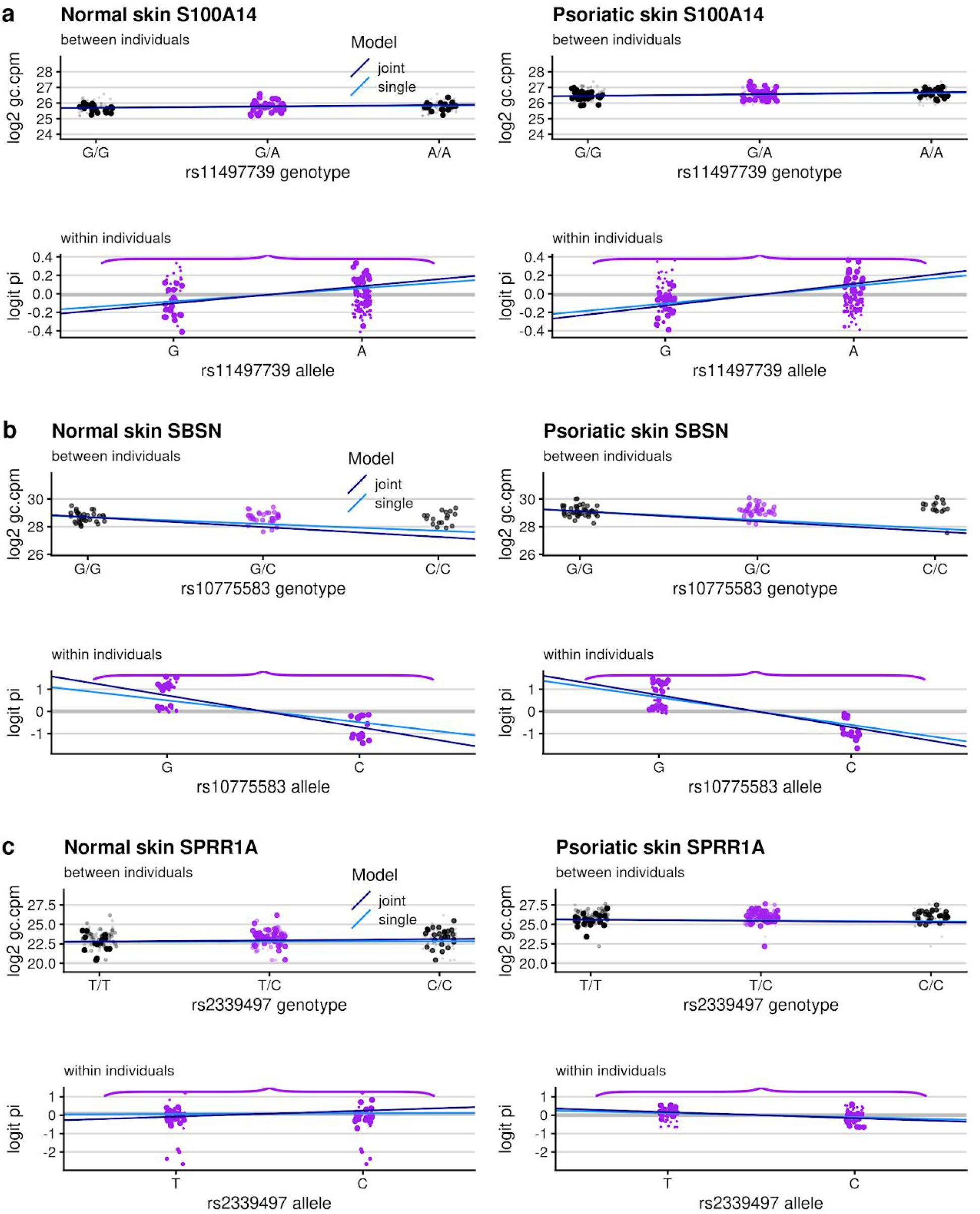
eQTL examples for the indicated genes. Same analysis as in Figure 6.

**Supplementary Figure 12.**
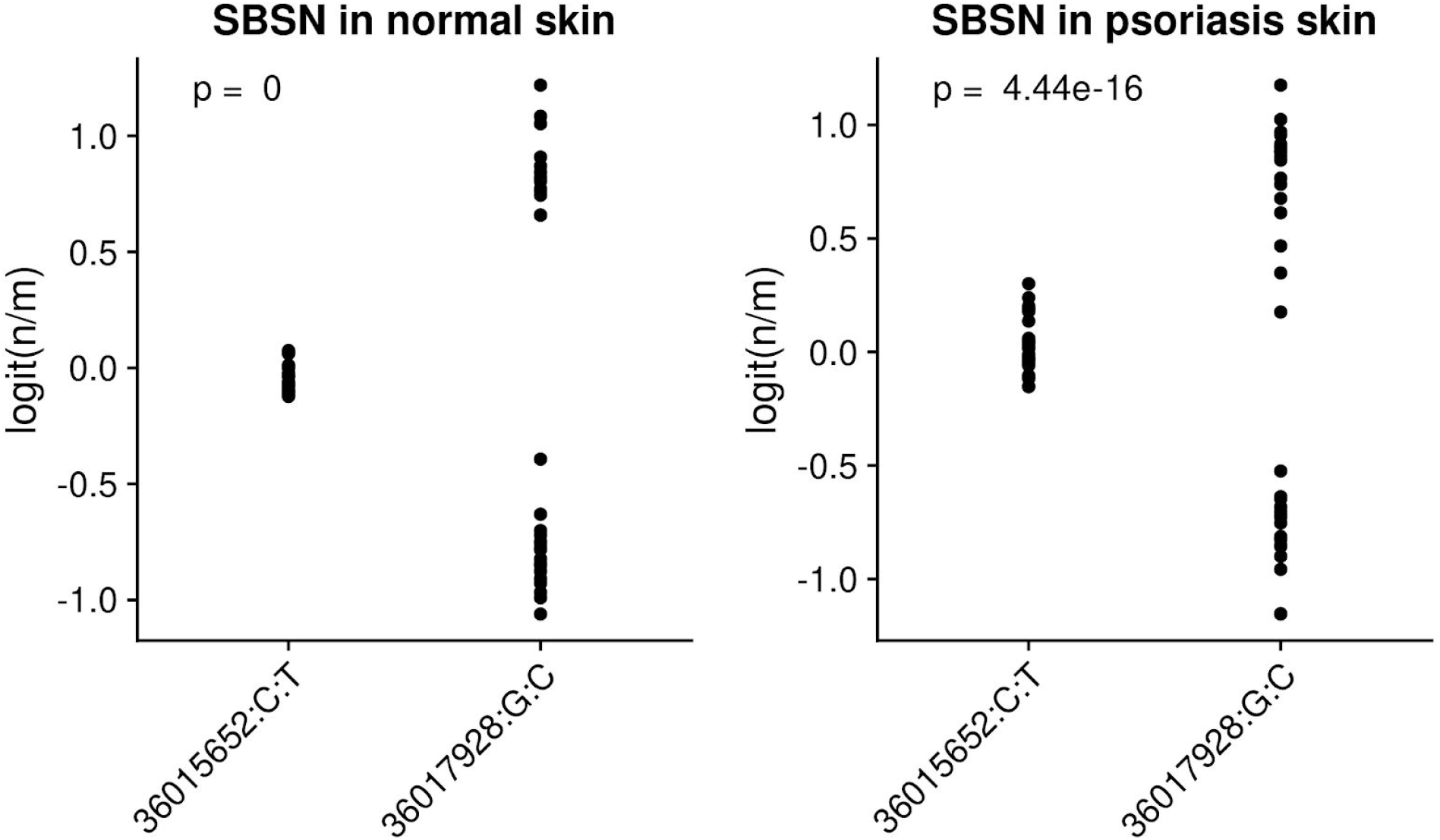
*SBSN* failed a post-hoc quality control measure testing homogeneity of variance of the raw allelic imbalance across fSNPs. Each plot corresponds to a skin type. For each fSNP (x-axis) the logit of the proportion of reads mapping the alternative allele for each individual (n/m) is shown in the y-axis. The p-value corresponds to a likelihood ratio test fitting beta binomial models allowing the overdispersion parameter to change across fSNPs or not.

**Supplementary Figure 13.**
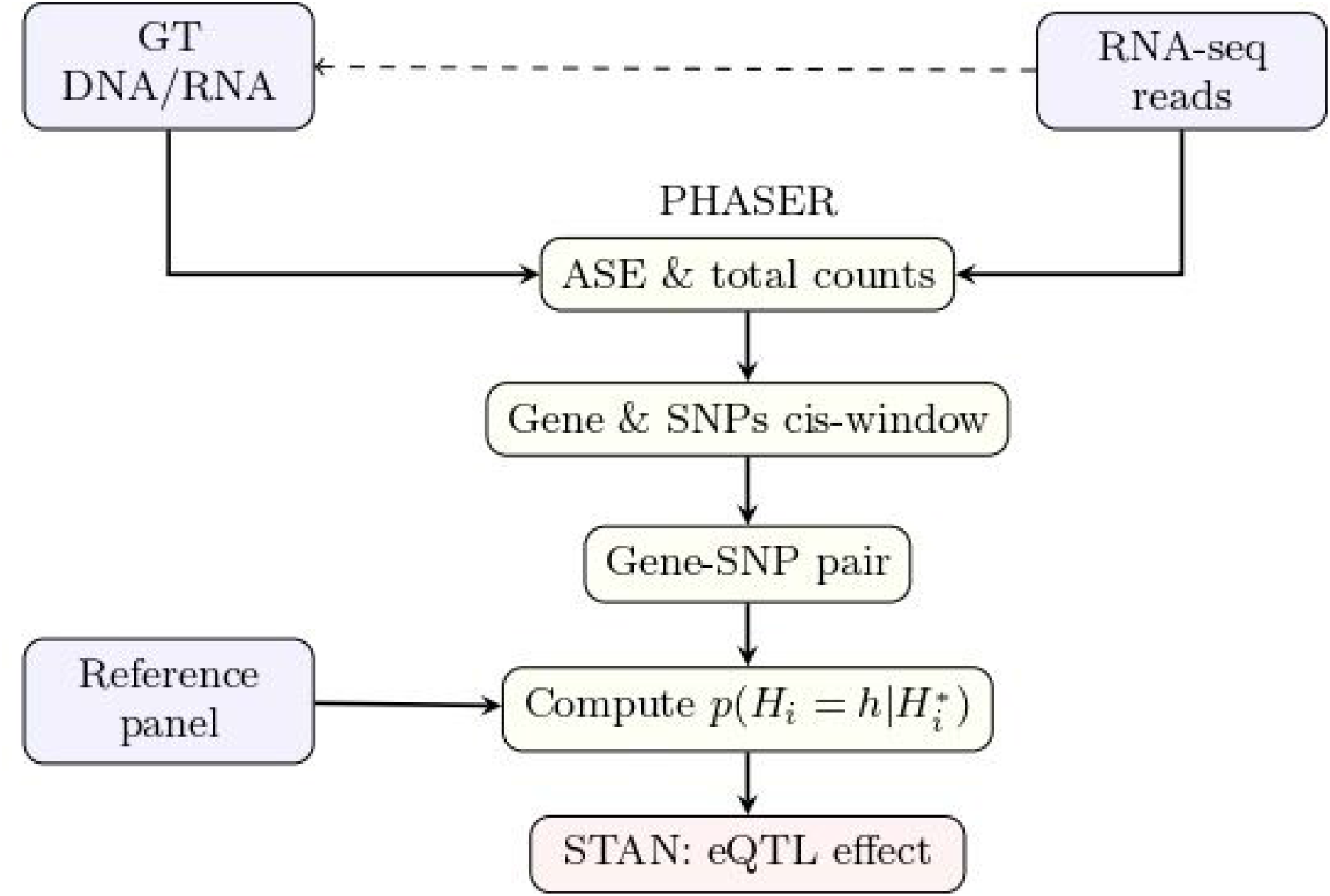
BaseQTL pipeline. Schematic diagram illustrating the different steps for input preparation and running BaseQTL.

**Supplementary Figure 14.**
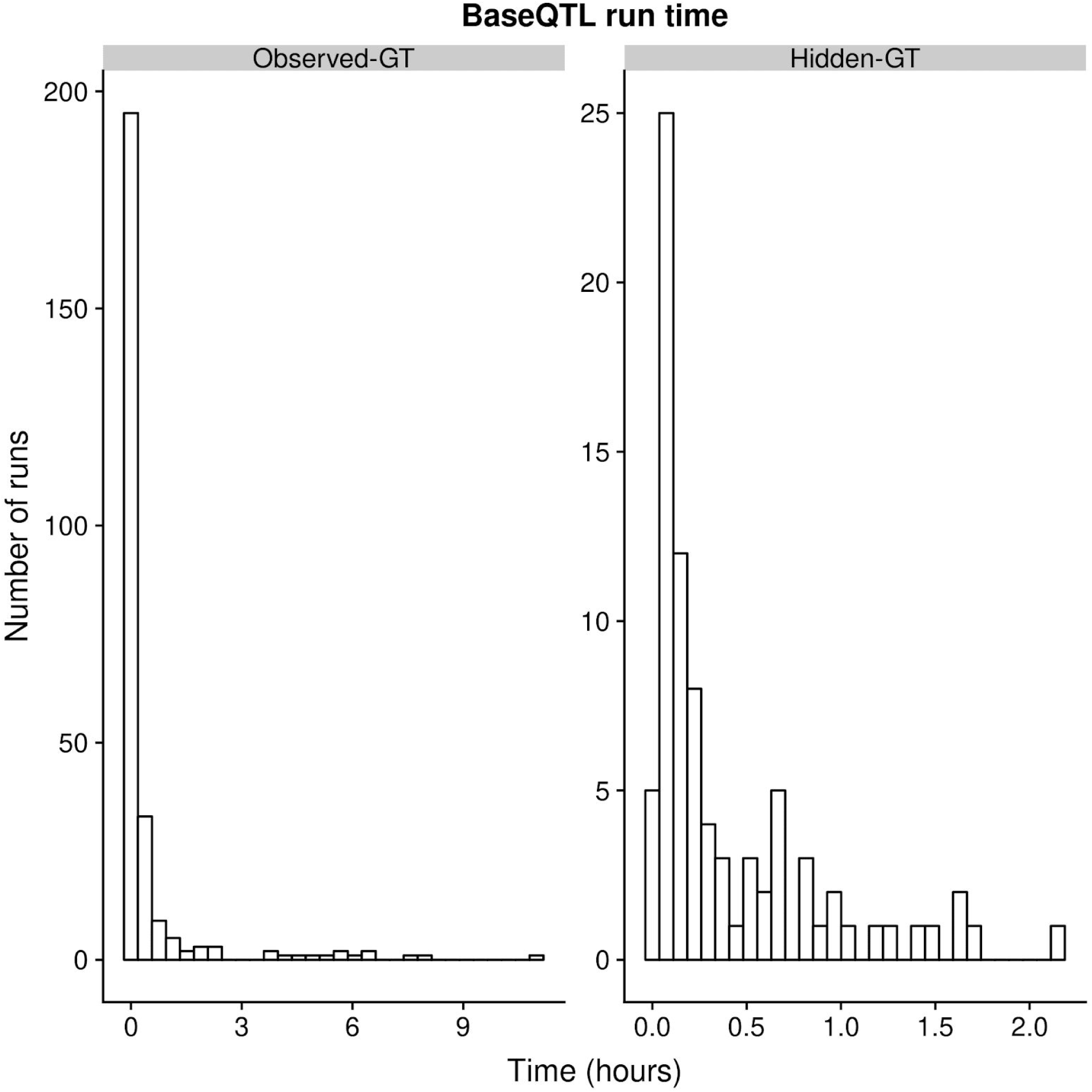
BaseQTL running time. The plot on the left shows the running time for the 264 genes run using the GEUVADIS dataset with observed genotypes, whereas the plot on the right corresponds to the 84 genes run with hidden genotypes. Each gene was run assessing candidate cis-SNPs within 1MB of gene using 16 cores. The median time was 6 and 10 minutes per gene for observed genotypes and hidden genotypes respectively, using a cis-window of 1MB.

**Supplementary Table 1:**
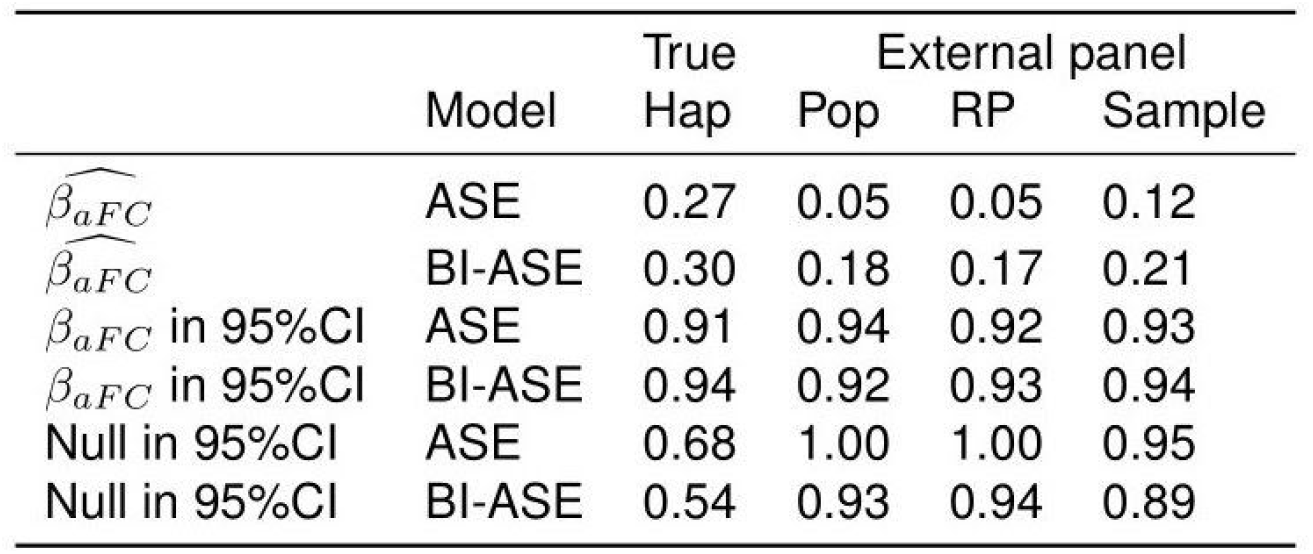
Effect of external panel on eQTL estimates. A population (Pop) of 50,000 haplotypes of a cis-eQTL and 3 fSNPs were simulated, with a *β*_*aFC*_ = 0.4 (Online Methods). From this population a random sample of 1000 haplotypes was extracted as was used as reference panel (RP). Samples of 100 haplotypes were also extracted from the population of haplotypes if the sum of the square difference of haplotype frequencies between the population and the sample relative to the haplotype frequency on the population was equal or higher than 0.05, for those haplotypes with frequency above 0.1 in the population. This procedure was repeated 100 times. For each of the 100 samples eQTL effects were estimated either with the full model (modelling both between-individual (Bl) and ASE signals) or ASE signals only. Each model was run with either known sample haplotypes (True haplotypes) or treating phasing as latent as estimating sample haplotypes using haplotypes from the population, the reference panel or the sample itself. The table shows the mean 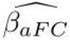 the proportion of times *β*_*aFC*_ is in the 95% credible interval and the proportion of times than the null value is within the 95% credible interval.This shows that inaccurate haplotype frequencies in the reference panel may lose power (the 95% Cl more often contains the null), but does not cause bias (the 95% Cl has 95% coverage of the true *β*_*aFC*_.

**Supplementary Table 2:** Gaussian components identified from fitting a mixture models (Online Methods) to GTEx eQTL estimates for the indicated tissues.

| Tissue | sd1 | weight1 | sd2 | weight2 |
| --- | --- | --- | --- | --- |
| Blood | 0.03 | 98% | 0.30 | 2% |
| LCL | 0.03 | 97% | 0.35 | 3% |
| Skin | 0.03 | 97% | 0.33 | 3% |

**Supplementary Table 3:** Error rates and missing genotypes for genotyping by RNA-seq. For the fSNPs used in inference with hidden genotypes, the number of erroneous calls and missing genotypes across all samples is shown.

| Errors | N | % | % (excluding missing) |
| --- | --- | --- | --- |
| 0 | 17151 | 72.00 | 99.32 |
| 1 | 116 | 0.49 | 0.67 |
| 2 | 1 | 0.00 | 0.01 |
| Missing | 6554 | 27.51 |  |

